# New tools for evaluating protein tyrosine sulphation: Tyrosyl Protein Sulphotransferases (TPSTs) are novel targets for RAF protein kinase inhibitors

**DOI:** 10.1101/296707

**Authors:** Dominic P Byrne, Yong Li, Pawin Ngamlert, Krithika Ramakrishnan, Claire E Eyers, Carrow Wells, David H Drewry, William J Zuercher, Neil G Berry, David G Fernig, Patrick A Eyers

## Abstract

Protein tyrosine sulphation is a post-translational modification (PTM) best known for regulating extracellular protein-protein interactions. Tyrosine sulphation is catalysed by two Golgi-resident enzymes termed Tyrosyl Protein Sulpho Transferases (TPSTs) 1 and 2, which transfer sulphate from the co-factor PAPS (3’-phosphoadenosine 5’-phosphosulphate) to a context-dependent tyrosine in a protein substrate. A lack of quantitative tyrosine sulphation assays has hampered the development of chemical biology approaches for the identification of small molecule inhibitors of tyrosine sulphation. In this paper, we describe the development of a non-radioactive mobility-based enzymatic assay for TPST1 and TPST2, through which the tyrosine sulphation of synthetic fluorescent peptides can be rapidly quantified. We exploit ligand binding and inhibitor screens to uncover a susceptibility of TPST1 and 2 to different classes of small molecules, including the anti-angiogenic compound suramin and the kinase inhibitor rottlerin. By screening the Published Kinase Inhibitor Set (PKIS), we identified oxindole-based inhibitors of the Ser/Thr kinase RAF as low micromolar inhibitors of TPST1/2. Interestingly, unrelated RAF inhibitors, exemplified by the dual BRAF/VEGFR2 inhibitor RAF265, were also TPST inhibitors *in vitro*. We propose that target-validated protein kinase inhibitors could be repurposed, or redesigned, as more-specific TPST inhibitors to help evaluate the sulphotyrosyl proteome. Finally, we speculate that mechanistic inhibition of cellular tyrosine sulphation might be relevant to some of the phenotypes observed in cells exposed to anionic TPST ligands and RAF protein kinase inhibitors.

**SUMMARY STATEMENT:** We develop new assays to quantify tyrosine sulphation by the human tyrosine sulphotransferases TPST1 and 2. TPST1 and 2 catalytic activities are inhibited by protein kinase inhibitors, suggesting new starting points to synthesise (or repurpose) small molecule compounds to evaluate biological TPST using chemical biology.

## INTRODUCTION

Like tyrosine phosphorylation [1], reversible tyrosine sulphation is a critical covalent modification that occurs on proteins post-translationally [2]. Originally identified more than half a century ago in sulphated fibrinogen and gastrin [3], tyrosine sulphation occurs on a wide range of secreted polypeptides in multicellular eukaryotes, and constitutes the transfer of a negatively charged sulphate group from the sulphate donor PAPS (3′-phosphoadenosine-5′-phosphosulphate) to a phenolic tyrosine residue. Tyrosine sulphation is catalyzed by two Golgi-associated membrane enzymes termed Tyrosyl Protein Sulpho Transferase 1 and 2 (TPST1 and 2), and sulphation leads to biologically-relevant changes in a large number of protein activities [2]. For example, sulphation can change the affinity of extracellular protein-protein interactions, such as those involved in chemotaxis [4] and host-pathogen interactions [5]. It also controls the proteolytic processing of both bioactive peptides [6, 7] and secreted antibodies [8], and multi-site tyrosine sulphation can change the function of several blood-coagulation regulators, including factor VIII [9, 10]. Interest in the pathophysiological analysis and therapeutic targeting of tyrosine sulphation was heightened by the finding that N-terminal chemokine receptor tyrosine sulphation in the HIV G-protein coupled receptor CCR5 [11, 12] plays a crucial role in coat binding and viral infection. Earlier studies had implicated tyrosine sulphation in the proteolytic control of the complement cascade component through decreased activity of C4 [13], the generation of gastrin from progastrin [14], and in regulating the binding of amino terminal sulphated P-selectin glycoprotein ligand-1 (PSGL-1) to P-selectin [15]. Interestingly, the binding of L-selectin on lymphocytes to mucin-like glycoproteins on endothelial cells is also regulated by sulphation, although the sialyl LewisX surface antigen is modified by a distinct carbohydrate 6-O sulphotransferase [16].

TPST1 was originally purified from bovine adrenal medulla [17, 18], and distinct human TPST1 and TPST2 genes have been cloned [19], with expression patterns varying markedly in both cells and tissues [20-22]. Both enzymes are believed to reside in the trans-Golgi compartment of the secretory pathway, and as type II transmembrane-containing enzymes with >85% sequence similarity in the intracellular catalytic domains, which are luminal-facing for substrate modification [19, 21, 22]. TPSTs interact with the sulphate-donor cofactor PAPS and an appropriate (often acidic) tyrosine-containing protein substrate. Recent experiments suggest that TPST1 and TPST2 might function as homo or heterodimers [23, 24], providing regulatory opportunities for the control of site-specific sulphation amongst substrates. In general, tyrosine sulphation occurs in an acidic context in proteins and model substrates [2, 18, 24-26], although some, including the bioactive protein gastrin, lack acid residues adjacent to the site of sulphation [14]. Analysis of a variety of synthetic peptides and intact proteins confirms that TPST1 and TPST2 can also control site-specific sulphation on multiple tyrosine residues, which are often clustered, consistent with a processive mechanism of modification [7, 27], or directionally distributed towards the substrate N-terminus [20, 28]. Crystal structures of TPST1 complexed with substrate peptides that are sulphated with different efficiencies have also been reported, and comparative analysis suggests differential substrate preferences for acidic residues adjacent to the site of modification [24, 29]. Structural comparison suggests a shared catalytic mechanism and substrate-binding energetics, driven by charge-based dynamic interactions. Bioinformatics analysis hints at a substantial and complex tyrosine sulphoproteome [30, 31] so uncovering the extent, substrate determinants and biological function of tyrosine-sulphated proteins remains a high priority technical challenge for Mass Spectrometry (MS)-based proteomics [32].

The analysis of tyrosine sulphation currently relies heavily on genetic and relatively low-throughput MS-based analysis, and only a few low-affinity inhibitors of TPSTs have ever been reported. Moreover, due to a lack of chemical tool compounds, biological sulphation remains highly understudied in general, relying on non-specific cytotoxic compounds such as chlorate with which to induce non-specific effects on sulphation [33]. The similarity between the sulphotransferase co-factor PAPS, and the phosphate donor ATP (utilised by protein kinases) raises questions as to whether PAPS-dependent sulphotransferases might be broad inhibitory targets for new or repurposed small molecules that target nucleotide-binding sites, especially well-studied families of compounds such as protein kinase inhibitors. Moreover, the mode of substrate peptide recognition observed in substrate and co-factor bound TPST2 structures closely resembles that established for the insulin-receptor tyrosine kinase bound to a tyrosine-containing (YMXM) substrate and ATP analogue [34], inviting further comparison between TPSTs and the highly druggable protein kinase superfamily [35, 36].

Analysis of TPST-based catalysis using small molecules remains in its infancy, and is currently hampered by a lack of rapid, flexible and reliable assays with which to screen for suitable inhibitors. Conventional procedures employ ^35^S-based detection of sulphated tyrosine in synthetic peptides [18, 21, 37] or, increasingly, rely on gas phase Mass Spectrometric (MS)-based detection of sulphated peptides [38-40]. Both of these approaches have technical drawbacks, and can be time-consuming, although ^35^S-based peptide sulphation by TPST2 was used to discover the first low-affinity reversible TPST2 inhibitors from a combinatorial library of aldehyde-linked heterocyclic compounds [37]. Recently, indirect fluorescent assays have been reported, including a PAPS depletion/reconstitution approach to monitor sulphate transfer [28] and continuous TPST1 and 2 assays reporting fluorescence-induced peptide sulphation [40]. The latter approach monitors peptide sulphation over relatively long periods of time, and requires inflexible positioning of the fluorophore relative to the modified tyrosine and flanking amino acid sulphation determinants. Nonetheless, such assays can be employed to discover small molecule inhibitors in screens, with several anionic compounds recently identified and cross-validated from commercial libraries [40, 41].

In this paper, we describe novel differential scanning fluorimetry (DSF) and sulphation assays that permit real time analysis of TPST1 and TPST2-mediated peptide sulphation, allowing us to evaluate TPST interactions with a variety of ligands and small molecule inhibitors. PAPS-dependent sulphation of peptides leads to a charge-induced mobility change, driven through intrinsic properties of a sulphotyrosine-containing substrate. Sulphation is detected by real-time mobility shift using a fluorescent microfluidic assay originally developed for the detection of peptide tyrosine phosphorylation [42]. In conjunction with analytical DSF, we also screened kinase inhibitor libraries, identifying a variety of known ligands as new TPST1 and TPST2 inhibitors, including the promiscuous protein kinase inhibitor rottlerin and a family of oxindole-based RAF kinase inhibitors from the Published Kinase Inhibitor Set (PKIS). In a related paper, published back-to-back with this study, we demonstrate that some of these compounds also inhibit the oligosaccharide sulphotransferase activity of Heparan Sulphate 2-O transferase (HS2ST), a related PAPS-dependent enzyme. Finally, chemically distinct inhibitors with activity towards the proto-oncogenic kinase RAF, exemplified by the dual BRAF/VEGFR2 inhibitor RAF265 (CHIR-265), were discovered to be more specific TPST inhibitors *in vitro*. We propose that target-validated kinase inhibitors might be chemically repurposed, or redesigned, to create new classes of TPST inhibitor. Moreover, we speculate that inhibition of cellular tyrosine sulphation by some of these compounds might contribute to the phenotypes observed in cells exposed to RAF kinase inhibitors.

## EXPERIMENTAL

### MATERIALS AND METHODS

#### Chemicals and Compounds

All standard biochemicals were purchased from either Melford or Sigma, and were of the highest analytical quality that could be obtained. PAPS (adenosine 3’-phosphate 5’-phosphosulphate, lithium salt hydrate), APS (Adenosine 5’-phosphosulphate, sodium salt), PAP (adenosine 3’-5’-diphosphate, disodium salt), CoA (coenzymeA, sodium salt) dephosphoCoA (3’-dephosphoCoA, sodium salt hydrate), ATP (adenosine 5’-triphosphate, disodium salt hydrate) ADP (adenosine 5’-diphosphate, disodium salt), AMP (adenosine 5’-monophosphate, sodium salt), GTP (guanosine 5’-triphosphate, sodium salt hydrate), GDP (guanosine 5’-diphosphate, sodium salt) or cAMP (adenosine 3’,5’-cyclic monophosphate, sodium salt) were all purchased from Sigma and stored at -80°C to ensure maximal stability. Rottlerin, surmain, aurintricarboxylic acid and all named kinase inhibitors were purchased either from Sigma, BD laboratories, Selleck or Tocris.

#### Cloning, protein purification and protein analysis

Human TPST1 (residues Lys43-Leu360) and TPST2 (residues Gly43-Leu359) enzymes lacking the transmembrane domains were amplified by PCR and cloned into pOPINF (OPPF-UK) to produce recombinant protein containing an N-terminal 6xHis tag and a 3C protease cleavage site. Recombinant TPST proteins were expressed in BL21 (DE3) pLysS *E. coli* (Novagen) with 0.4 mM IPTG for 18 h at 37°C and isolated from inclusion bodies and refolded as previously described [43]. In brief, cells were resuspended in 3 ml ice-cold lysis buffer (50 mM Tris-HCl, pH 8.0; 10 mM MgCl_2_; and 1 mM DTT supplemented with cOmplete, EDTA-free protease inhibitor cocktail tablets (Roche) per gram of *E. coli* cell pellet, and flash frozen with liquid nitrogen. Cells were disrupted by sonication, inclusion bodies were collected by centrifugation for 1 h at 10,000 x g at 4 °C and washed in ice-cold WB1 (50 mM Tris-HCl, pH 8.0; 100 mM NaCl; 10 mM EDTA and 1 % (v/v) Triton X-100) followed by WB2 (20 mM Tris-HCl, pH 8.0; 200 mM NaCl and 1 mM EDTA). Inclusion bodies were resuspended in SB (100 mM Tris-HCl, pH 8.0; 6 M GndHCl; 5 mM EDTA and 10 mM DTT) and incubated at 4 °C with constant agitation. SB buffer was supplemented with fresh DTT (10 mM DTT) after 12 h and incubated for 2 h at room temperature. Insoluble material was removed by centrifugation (1 h, 60,000 x g, 4 °C), and the supernatant was concentrated by ultrafiltration (Amicon Ultra-15 centrifugal filter unit, 10 kDa cutoff) and then diluted 10 fold with buffer A (100 mM Na acetate, pH 4.5; 6 M GndHCl and 10 mM DTT). 5 ml of concentrated TPST (~150 mg) was slowly added (using a peristaltic pump) to 1 L of pre-chilled refolding buffer (50 mM Tris-HCl, pH 8.5; 500 mM GndHCl; 10 mM NaCl; 0.4 mM KCl; 0.1 mM EDTA; 0.14 mM DDM; 5 mM GSG and 2.5 mM GSSG) while mixing with a magnetic stirrer. The refolding mixture was incubated for 20 h without mixing at 4 °C and precipitated protein was removed by centrifugation. Soluble TPST protein was then purified by immobilized metal affinity chromatography and size-exclusion chromatography (SEC) using a HiLoad 16/600 Superdex 200 column (GE Healthcare) equilibrated in 50 mM Tris– HCl, pH 7.4, 100 mM NaCl, and 10% (v/v) glycerol. Glutathione-*S*-transferase (GST) tagged CC4 tide (EDFEDYEFDG) was cloned into pOPINJ (OPPF-UK) and affinity purified from BL21 (DE3) pLysS *E. coli* using Glutathione-Sepharose 4B (GE Healthcare) and size-exclusion chromatography. The tyrosine kinase EphA3, comprising the kinase domain and the juxtamembrane region with an N-terminal 6xHis-tag, was expressed in pLysS *E. coli* from pET28a LIC, and protein purified using Ni NTA agarose and gel filtration, as described [42]. Halo-FGF7 was purified as previously described [44].

#### SDS-PAGE and immunoblotting

After assay, proteins were denatured in Laemmli sample buffer, heated at 95 °C for 5 min and then analysed by SDS-PAGE with 10% (v/v) polyacrylamide gels. Gels were stained and destained using a standard Coomassie Brilliant Blue protocol. To evaluate protein sulphation and phosphorylation by immunoblotting, standard western blotting procedures were followed using an anti-sulphotyrosine antibody (Millipore) in the presence of appropriate positive and negative controls, and modifications visualised using ECL reagent.

#### DSF assays

Thermal shift/stability assays (TSAs) were performed with a StepOnePlus Real-Time PCR machine (Life Technologies) using Sypro-Orange dye (Invitrogen) and thermal ramping between 20 - 95°C in 0.3°C step intervals per data point to induce denaturation in the presence or absence of various biochemical and small molecule inhibitors [45, 46]. TPST1 and TPST2 were assayed at a final concentration of 5 μM in 50 mM Tris–HCl (pH 7.4) and 100 mM NaCl. Final DMSO concentration in the presence or absence of the indicated concentrations of ligand was no higher than 4% (v/v). None of the test compounds analysed in the absence of HS2ST were found to interfere with fluorescent detection of Sypro-Orange binding. Normalized data were processed using the Boltzmann equation to generate sigmoidal denaturation curves, and average *T*_m_/Δ*T*_m_ values were calculated as described [47] using GraphPad Prism software.

#### EZ Reader II-based peptide sulphation assays

Fluorescently-tagged peptides used in TPST sulphotransferase assays were derived from the human physiological substrate sequences where noted. A 5-FAM fluorophore, with maximal absorbance of 495 nm and a maximal emission absorbance of 520 nm that could be detected in an EZ Reader *via* LED-induced fluorescence, was covalently coupled to the free N-terminus of each peptide. CC4-tide (5-FAM-EDFEDYEFDG-CONH_2_ and the equivalent peptide lacking the single acceptor tyrosine residue, 5-FAM-EDFEDFEFDG-CONH_2_), were modified from the human Complement C4 protein [13], Fibroblast Growth Factor 7 (FGF7, 5-FAM-ERHTRSYDYMEGGD-CONH_2_), C-C motif chemokine receptor 8 (CCR8, 5-FAM-TTVTDYYYPDIFSS-CONH_2_) and P-selectin glycoprotein ligand-1 (PSGL1, 5-FAM-TEYEYLDYDFLPETE-CONH_2_) peptides were derived from the appropriate human sequences (predicted site of tyrosine sulphation shaded in red). Peptides were synthesised using solid-phase Fmoc chemistry and after HPLC purification (>95%), the expected intact peptide mass was confirmed by MALDI-TOF Mass Spectrometry (Pepceuticals, Leicester, UK). The Perkin Elmer LabChip EZ II Reader system [48], 12-sipper chip and CR8 coating, assay separation buffer and a synthetic fluorescent Ephrin A3 substrate (Ephrin A3-tide, 5-FAM EFPIYDLPAKK-CONH_2_) were all purchased from Perkin Elmer. Pressure and voltage settings were adjusted manually to afford optimal separation of tyrosine sulphated and non-sulphated peptides. Individual sulphation assays were performed in a 384 well plate in a volume of 80 µl in the presence of the indicated concentration of PAPS (Sigma-Aldrich), 50 mM HEPES, 0.015 % (v/v) Brij-35 and 5 mM MgCl_2_ (unless specified otherwise) and the degree of peptide sulphation was directly calculated by EZ Reader software by differentiating sulphopeptide:peptide ratios. The activity of TPST proteins in the presence of inhibitors was quantified by monitoring the amount of sulphopeptide generated over the assay time relative to control assay with no additional inhibitor molecule. Data was normalized with respect to these control assays, with sulphate incorporation into the peptide limited to ~ 20 % to prevent depletion of PAPS and to ensure assay linearity. K*m* and IC_50_ values were determined by non-linear regression analysis using Graphpad Prism software

#### Biochemical and small molecule screening by DSF and TPST enzyme assay

The PKIS chemical library (designated with SB, GSK or GW prefixes) comprises 367 largely ATP-competitive kinase inhibitors, covering 31 chemotypes originally knowingly designed to inhibit 24 distinct protein kinases [49, 50], was stored frozen as a 10 mM stock in DMSO at -80°C. This inhibitor library is characterised as highly drug-like (~70% with molecular weight <500 Da and clogP values <5). For initial screening, compounds pre-dissolved in DMSO were pre-incubated with TPST1 or TPST2 for 10 minutes and sulphotransferase reactions initiated by the addition of the universal sulphate donor PAPS. For inhibition assays, competition assays, or individual IC_50_ value determination, the appropriate compound range was prepared by serial dilution in the appropriate solvent, and added directly into the assay to the indicated final concentration. All control experiments contained 4% (v/v) DMSO.

#### Molecular docking analysis

Rottlerin, GW305074X, suramin and RAF265 were built using Spartan16 (https://www.wavefun.com) and energy minimised using the Merck molecular forcefield. GOLD 5.2 (CCDC Software;) was used to dock molecules [51], with the binding site defined as 10 Å around the 5’ phosphorous atom of PAP, using coordinates from human TPST1 PDB ID: 5WRI [24]. A generic algorithm with ChemPLP as the fitness function [52] was used to generate 10 binding-modes per ligand in HS2ST. Protons were added to the protein. Default settings were retained for the “ligand flexibility” and “fitness and search options”, however GA settings were changed manually to 200%.

## RESULTS

### Analysis of human TPST1 and TPST2 using a reliable thermal stability assay (TSA)

To drive the development of new approaches to assay and inhibit protein tyrosine sulphation, we developed a Differential Scanning Fluorimetry (DSF) assay to examine the thermal stability of TPST1 or TPST2 in the presence or absence of biochemical ligands (Figure 1A). We purified recombinant soluble human 6His-tagged TPST1 and 2 catalytic domains (amino acids 43-360 and 43-559 respectively, lacking the transmembrane domain) from bacterial inclusion bodies to near homogeneity (Figure 1B). After refolding from guanidine hydrochloride into a Tris-based buffer, TPST thermal stability and unfolding profiles were measured in the presence of the known sulphated co-factor PAPS, or the dephosphorylated precursor APS, whose phosphorylation at the 3’ position on the adenine ring by APS kinase generates PAPS in cells. Heating of TPST1 and TPST2 generated a typical heat-induced unfolding profile with both TPST1 and TPST2 exhibiting almost identical T_m_ values (formally, the temperature at which 50% of the protein is unfolded based on fluorescence) of ~40 °C (Figure 1C, E). In both cases, inclusion of PAPS in the unfolding assay induced a shift in the T_m_ value, suggesting that both enzymes were folded and could bind to a physiological co-factor. In the case of TPST1, PAPS (but not APS) induced a ΔT_m_ value of ~3 °C (Figures 1C, D), whereas TPST2 stability shifted by ~9 °C in the presence of PAPS, but not APS (Figures 1E, F). Side-by-side comparison of TPST1 and TPST2 over a range of PAPS concentrations demonstrated concentration dependent effects on TPST stability, with a more marked shift in TPST2 stability at all concentrations tested (Figure 1G). We next compared thermal unfolding in the presence of a panel of nucleotide-based cofactors. These experiments demonstrated a lack of significant thermal shift by Mg^2+^ ions, APS, AMP or cAMP. In contrast, ADP, PAP, CoA, acetyl CoA and GTP all induced marked stabilisation of both TPST1 and TPST2 at near stoichiometric concentrations in the assay, suggestive of high affinity binding. In contrast to CoA, dephospho-CoA, which lacks a 3’-phosphoadenine group, was unable to induce thermal shifts in either TPST1 or 2, as established for APS, in which the 3’ phosphoadenine group is also absent. In the case of ATP, ADP and GTP, TPST1 and TPST2 shifts were abolished in the presence of Mg^2+^ ions, presumably reflecting the very high affinity of this divalent cation for these nucleotides [53] (Supplementary Figure 1A, B).

**Figure 1.**
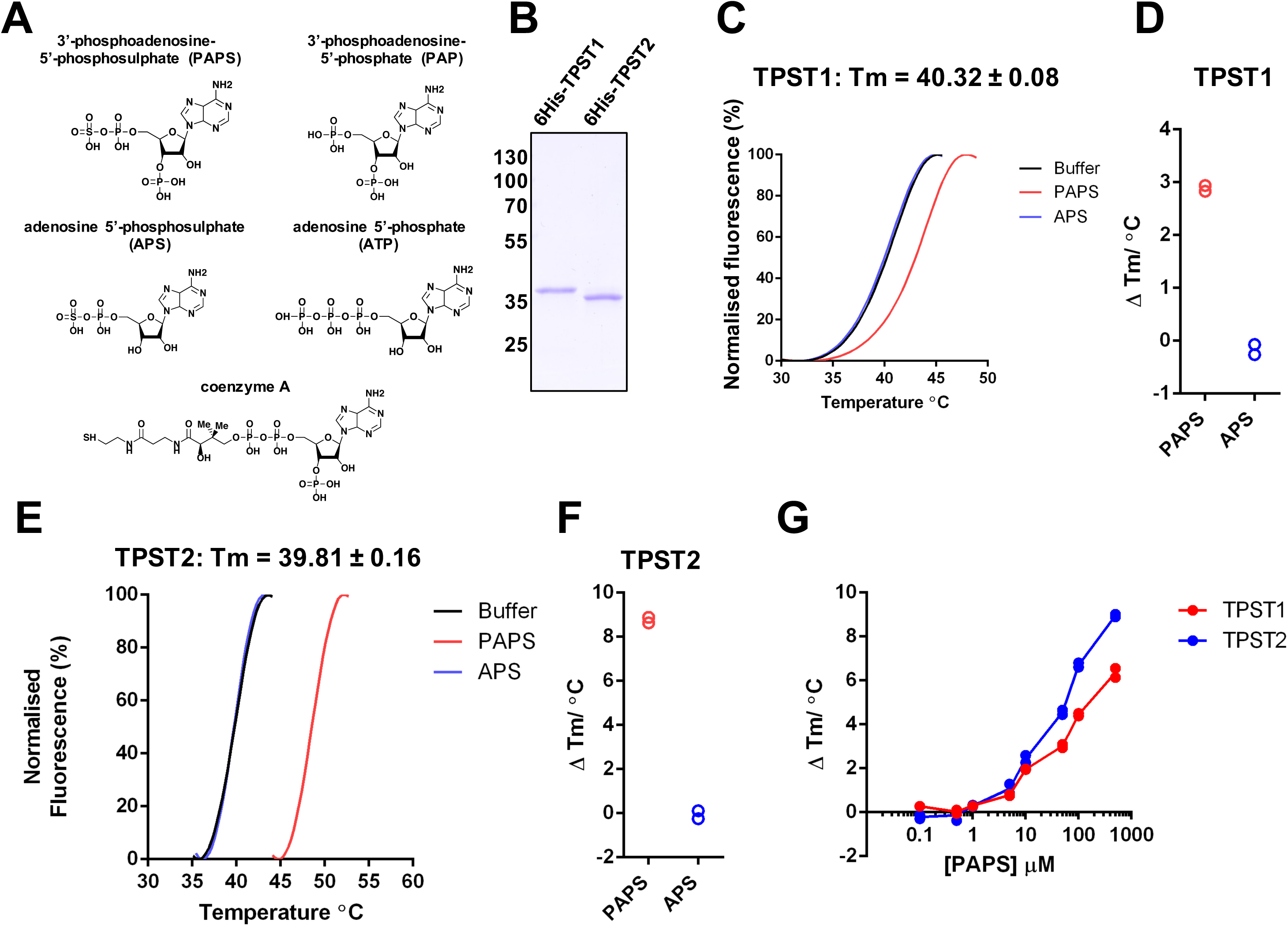
Analysis of purified recombinant 6His-TPST proteins. **(A)** Biochemical structure of PAPS and PAPS-related compounds. **(B)** Coomassie blue staining of purified recombinant 6His-TPST enzymes: 1 μg of TPST1 and 2 were analysed by SDS-PAGE after purification to near homogeneity. **(C)** TSA, and calculation of T_m_, for TPST1 (5 μM) in the presence of 0.5 mM PAPS (red) or 0.5 mM APS (blue); buffer control is in black. **(D)** ∆T_m_ for TPST1 in the presence of PAPS and APS, as measured by DSF, data derived from **(C)**. ∆T_m_ values were calculated by subtracting the control T_m_ value (buffer, no nucleotide) from the measured T_m_ value. **(E)** As for **(C)** but using TPST2. **(F)** As for **(D)**, but employing TPST2. **(G)** Analysis of PAPS-dependent thermal stabilization of TPST1 and TPST2. TSA employing TPST1 or TPST2 proteins (5 μM) were measured in the presence of the indicated concentration of PAPS. ∆T_m_ values were calculated by DSF, as described above.

### A novel microfluidic assay to quantify real-time peptide sulphation by TPST1 and TPST2

Thermal and enzymatic screening assays can generate complementary information to help evaluate ligand binding. [54] To extend our thermal analysis of TPST ligand binding to include real-time analysis of sulphate transfer, and help progress our eventual goal of discovering TPST1 and TPST2 inhibitors, we developed a novel enzyme assay for kinetic analysis of peptide tyrosine sulphation. The basic requirement of this assay was that it should report the enzymatic incorporation of sulphate onto a tyrosine residue of a synthetic peptide substrate with a high signal-to-noise ratio and be rapid, repeatable and relatively high-throughput. Current protocols to monitor tyrosine sulphation generally involve ^35^S-based enzyme regeneration or intrinsic fluorescence assays, which are often unsuitable for kinetic or high-throughput analysis using different peptide substrates, and are prone to artefacts if compounds or co-factors that interfere with fluorescence detection are employed. However, as established below, our novel assay permits rapid real-time detection of non-radioactive sulphate incorporation into synthetic peptides.

### Synthetic peptides derived from human substrates are *in vitro* TPST1 and/or TPST2 substrates

To evaluate context-specific sulphation kinetics for TPST1 and 2, we synthesised a panel of peptides possessing tyrosine-containing sequences found in human proteins previously reported to be sulphated on tyrosine [2], and developed an assay to quantify peptide sulphation. The assay comprises a putative substrate peptide (containing a tyrosine in an acid context and culminating in an amide group), TPST1 or TPST2 and the PAPS co-factor (Figure 2A). To facilitate the facile detection of both sulphated and non-sulphated substrates in the same assay using a microfluidic platform, we appended an N-terminal fluorophore (5-FAM) to the peptide. Since tyrosylsulphate (singly charged under the assay conditions) and tyrosylphosphate (doubly charged) are chemically similar, and can potentially induce charge based differences in peptide mobility when covalently attached to tyrosine, we reasoned that this assay would be able to detect sulphation in a similar way to that previously established for tyrosine phosphorylation by Ser/Thr and Tyr kinases [42, 55, 56]. As shown in Figure 2B, incubation of a 5-FAM conjugated 10-mer tyrosine-containing peptide from human complement C4 protein with TPSTs led to the appearance of an electrophoretically distinct product (P) when compared with the unmodified substrate (S). Different ratios of product to substrate were detected when TPST1 or TPST2 were included in the assay, but no product was detected with buffer and PAPS alone, suggesting that the new product was a tyrosine-sulphated peptide (Figure 2B).

**Figure 2.**
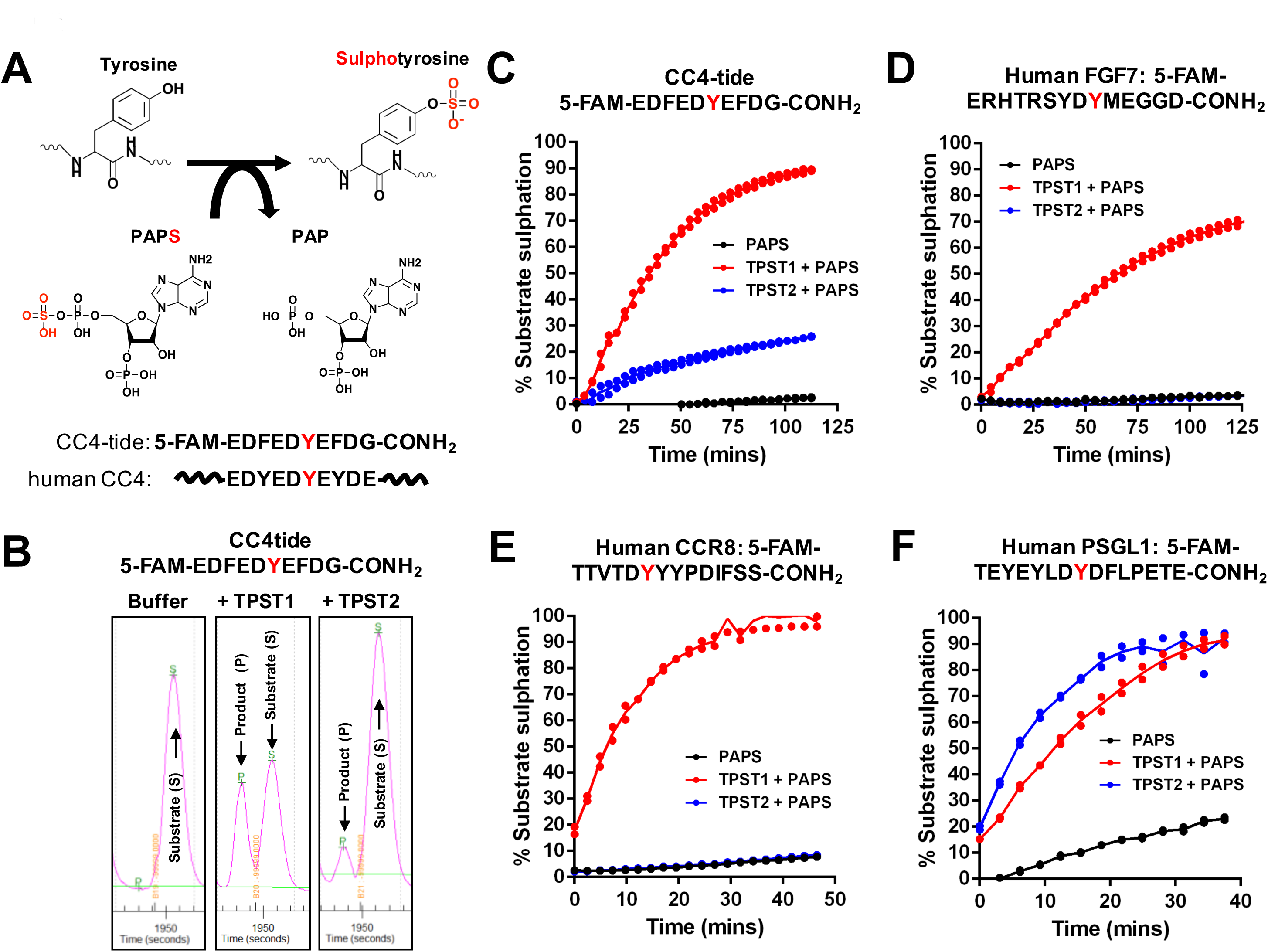
Detection of tyrosylprotein sulphotransferase activity using a direct microfluidic mobility shift assay and fluorescent peptide substrates. **(A)** Schematic representation of PAPS-dependent sulphate incorporation into a tyrosine residue of a substrate. The sequence of the synthetic single Tyr-containing peptide (CC4-tide, containing a fluorescent 5-FAM group at the N-terminus) is shown above the native human CC4 protein sequence. **(B)** TPST1 and TPST2-dependent tyrosine sulphation alters the microfluidic mobility of CC4-tide. Separation of the higher-mobility, sulphated (product, P) peptide from the lower-mobility (substrate, S) peptide occurs through a difference in their net charge. **(C-F)** Time-dependent tyrosine sulphation of CC4-tide **(C)** or fluorescently-labelled tyrosine-containing substrate peptides derived from human FGF7 **(D),** CCR4 **(E)** or PSGL1 **(F)** proteins. Direct peptide sulphation was calculated by measuring the ratio of substrate peptide to sulpho-peptide at the indicated time points after adding the PAPS co-factor. All assays were performed at room temperature (20 °C) using 2 μM final concentration of the appropriate fluorescent peptide substrate, 500 μM PAPS and 0.4 μM TPST1 or TPST2.

We next compared the ability of TPST1 and TPST2 to modify the CC4 peptide (termed hereafter CC4-tide) in a kinetic assay, monitoring the real-time appearance of the sulphated peptide by detecting the increase in product peak height in a duplicate assay format. As shown in Figure 2C, TPST1 was much more efficient at modifying CC4-tide, inducing near-stoichiometric modification after one hour. The rate of sulphation by TPST1, but not TPST2, was responsive to divalent cations, and could be increased 6-10 fold by including Mg^2+^ ions or Mn^2+^ ions in the buffer (Supplementary Figure 3), despite a lack of detectable Mg^2+^ binding to TPST1 (or TPST2) by DSF (Supplementary Figure 1A, B), consistent with previous studies [18, 28, 40]. In contrast, over the same time period and at the same concentration in the assay, purified TPST2 only sulphated CC4-tide to a stoichiometry of ~20%. We next assessed whether TPST1 or TPST2 sulphated a fluorescent 14-mer peptide derived from recombinant human FGF7, which contains a single known site of tyrosine sulphation corresponding to Tyr27 in the mature growth factor [57]. Using this new assay, we were unable to detect FGF7-tide sulphation by TPST2, although TPST1 induced ~70% peptide sulphation over the assay time-course (Figure 2D). Interestingly, CCR8-tide, which was derived from the human CCR8 sequence, was even more rapidly sulphated by TPST1 than FGF7-tide, although it was not modified noticeably by TPST2 (Figure 2E). In marked contrast, a distinct fluorescent 14-mer sequence derived from human PSGL1 was rapidly, and stoichiometrically sulphated by TPST1 and TPST2 (Figure 2F), confirming these enzymes possess overlapping, as well as distinct, substrate specificities *in vitro*. As a test of the suitability of our assay to derive a reported kinetic parameter, we employed an appropriate concentration of TPST1 and 2 so that the degree of peptide sulphation was linear over the time course of the reaction. Under these conditions, the K*m* value for PAPS in a TPST1-dependent CC4-tide assay was 6.6 ± 1.9 µM, (Supplementary Figure 2) consistent with previous literature reports of 2-5 µM for TPST1 [18, 25] or 12 µM for TPST2 obtained from transfected CHO cell medium [37] or ~5 µM for recombinant TPST2 isolated from baculovirus [40]

### Substrate and co-factor specificity for model TPST1 and TPST2 substrates

To investigate TPST site specificity, we confirmed that the single Tyr residue in CC4 represents the site of covalent modification identified in the mobility assay, by generating a peptide in which Tyr was substituted for a chemically analogous, but non-sulphatable, Phe residue. As shown in Figure 3A and B, PAPS-dependent CC4-tide sulphation likely occurs on Tyr, because the Phe-substituted peptide was not modified by incubation with TPST1 (or TPST2). To confirm sulphate and phosphate co-factor specificity in the assay, we evaluated sulphation and phosphorylation of the same CC4-tide substrate using either TPST1 or the tyrosine kinase Ephrin A3. Importantly, Ephrin A3 generated a modified (phosphorylated) CC4 product peptide in the presence of ATP, but not PAPS, whereas TPST1 generated a modified (sulphated) product peptide only in the presence of PAPS, but not ATP (Figure 3C). The electrophoretic mobility of sulphated (TPST1-generated) or phosphorylated (EphA3-generated) CC4 peptides relative to the non-modified peptide was very similar in our microfluidic assay conditions, consistent with similar physiochemical properties of anionic sulphotyrosine and phosphotyrosine formed in the assay. Finally, we confirmed the reported preference for tyrosine sulphation of peptide substrates in the context of an N-terminal acidic residues (which is targeted electrostatically to the cationic active site) by showing that a tyrosine kinase (TK) peptide substrate optimised for EphA3 phosphorylation was not modified by TPST1 or TPST2, presumably because it lacked an acidic residue at the -1 position relative to Tyr, although this did not prevent phosphorylation by EphA3 kinase in the presence of Mg-ATP (Figure 3D).

**Figure 3.**
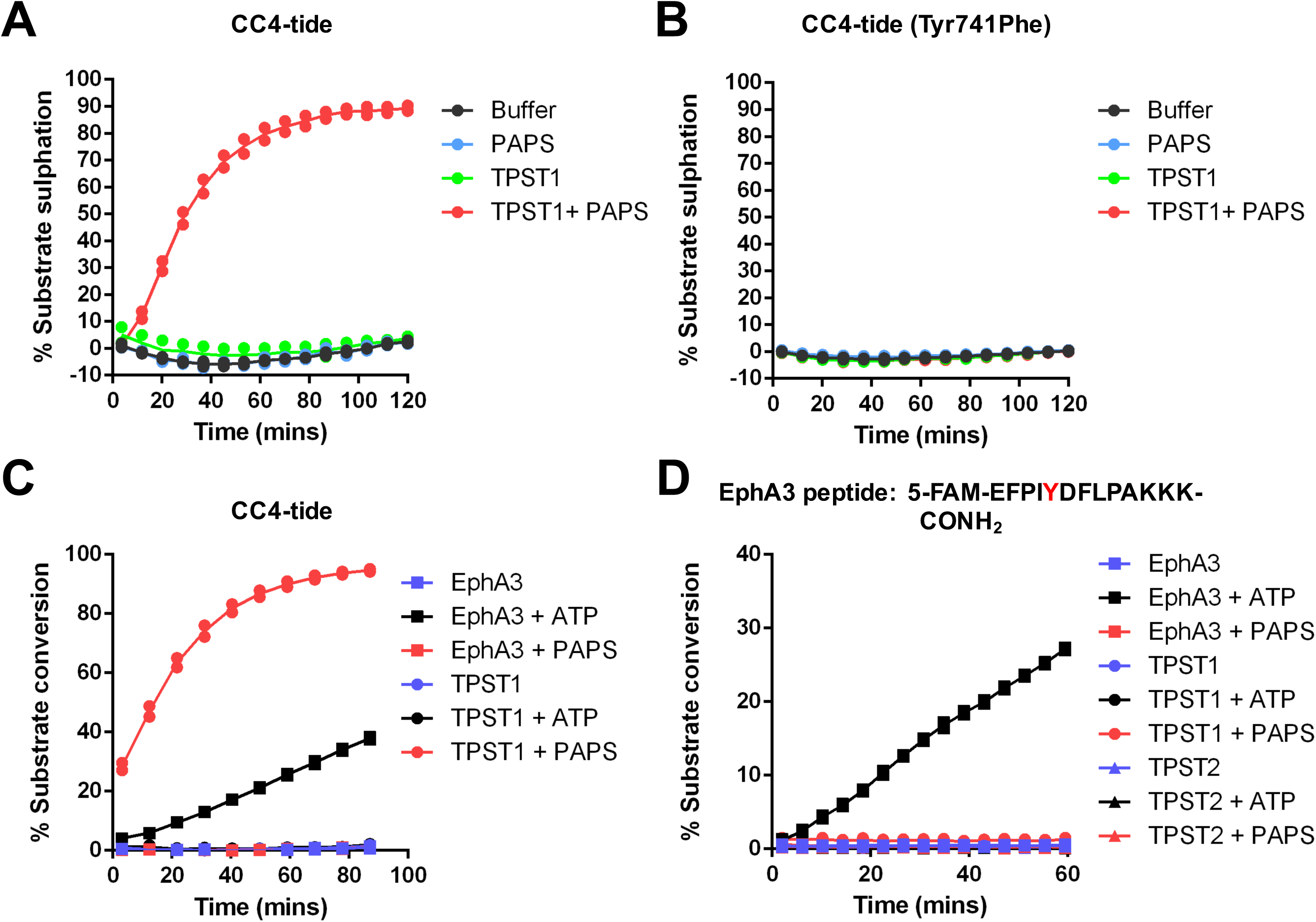
Changes in fluorescent peptide mobility are a consequence of TPST-catalysed tyrosine sulphation. Mobility-analysis of TPST1-dependent peptide sulphation for **(A)** CC4-tide or **(B)** CC4-tide in which the Tyr acceptor site is mutated to Phe (Tyr741Phe). Recombinant TPST1 enzyme (0.4 μM) was assayed using 2 μM peptide substrate ± 10 μM PAPS as sulphate donor. **(C)** Dual detection of tyrosine phosphorylation or tyrosine sulphation of CC4-tide. TPST1 (0.2 μM) or EphA3 tyrosine kinase (0.3 μM) were incubated with 2 μM CC4-tide in the presence of 500 μM PAPS (sulphate donor) or 500 μM ATP (phosphate donor). **(D)** Lack of tyrosine sulphation of a distinct tyrosine-containing EphA3 substrate peptide by TPST1 or 2. Assay conditions were as for **(C)**.

### TPST1 and TPST2 sulphate tyrosine in recombinant sulphoacceptor proteins

Detection of quantitative tyrosine sulphation using real-time microfluidics represents a totally new approach to study this covalent modification *in vitro*. In order to unambiguously confirm sulphation by TPST1 and TPST2 using a complementary technique, we used immunoblotting with a monoclonal antibody that specifically recognises sulphated tyrosine in intact proteins. Initially, we generated a recombinant protein consisting of Glutathione S-tranferase (GST) fused to the CC4-tide sequence (EDFEDYEFDG) that was developed for TPST1 and TPST2 mobility-based enzyme assays (Figures 2 and 3). As detailed in Figure 4A, this GST fusion protein became sulphated on tyrosine only after incubation with TPSTs, and this modification required PAPS in the assay. Consistently, GST was not detectably sulphated under any condition, confirming that the CC4-tide sequence was the target of both enzymes. Site specificity in the assay was confirmed by mutation of Tyr to Phe in the GST fusion protein, which abolished detection by the sulphotyrosine antibody. As a further control, we demonstrated that GST-CC4-tide could become Tyr phosphorylated, but not Tyr sulphated, after incubation with Ephrin A3 and Mg-ATP at the same tyrosine sulphated by TPST1/2 in GST-CC4-tide, (Figure 4C, note detection of pTyr in EphA3 protein due to autophosphorylation). This experiment also demonstrates unequivocally that the modification-specific antibodies can differentiate between sulphated and phosphorylated forms of the GST-CC4-tide protein. We also confirmed that full-length recombinant FGF7 was specifically modified by TPST1, but not TPST2 *in vitro*, consistent with side-by-side kinetic analysis of TPST1 and TPST2 FGF7-tide sulphation (compare Figures 2D and 4D).

**Figure 4.**
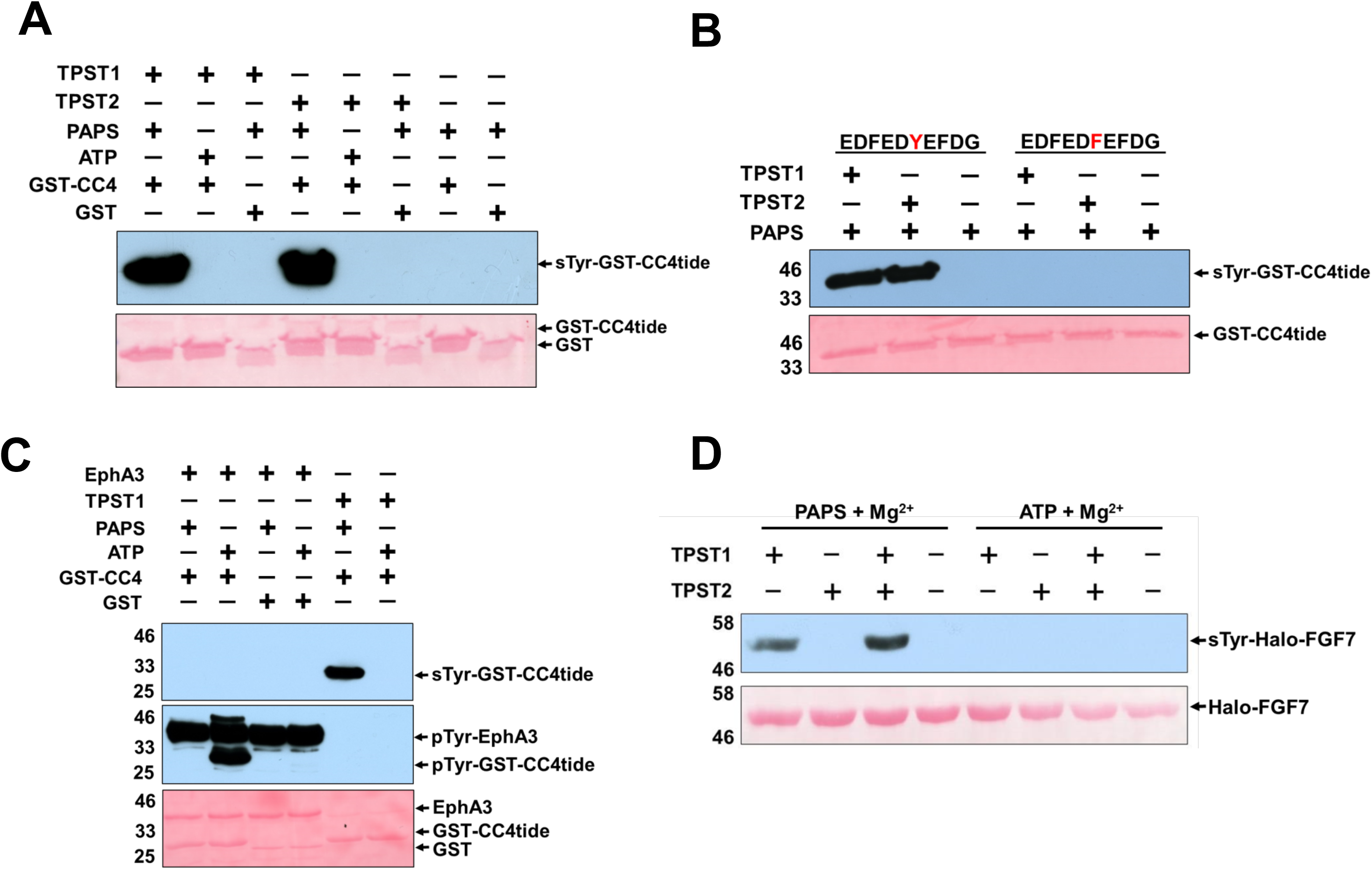
Validation of *in vitro* recombinant TPST sulphotransferase activity. Immunoblot of an *in vitro* sulphotransferase assay using a recombinant GST-tagged CC4-tide. Recombinant, purified GST-CC4-tide or purified GST alone (1 μg) were incubated at room temperature for 1 h with 1 μg TPST1 or TPST2 ± 500 μM PAPS (sulphate donor) or 500 μM ATP (phosphate donor). **(B)** Immunoblot demonstrating TPST1 and TPST2-dependent sulphation of GST-CC4-tide at the specific tyrosine residue sulphated in intact CC4 (Tyr741). Assay was performed with 1 μg of each TPST enzyme, GST-CC4tide or GST-CC4-tide (Tyr741Phe) and 500 μM PAPS for 1 h at room temperature. **(C)** Western blot confirming that monoclonal sulphotyrosine antibody does not cross react with phosphotyrosine. 1 μg GST-CC4-tide and GST was incubated for 1 h with 2 μg TPST1 or EphA3 ± 500 μM PAPS or 500 μM ATP. GST CC4-tide sulphation (top panel), EphA3 autophosphorylation or GST-CC4-tide phosphorylation (middle panel) are indicated. **(D)** Detection of recombinant FGF7 *in vitro* sulphation by immunoblot. Halo-tagged FGF7 (5 μg) was incubated for 16 h at 20°C with 2 μg TPST protein ± 500 μM PAPS 500 μM ATP and tyrosine sulphation was detected using a monoclonal sulphotyrosine antibody (top panel). For all assays, equal loading of substrate proteins was confirmed by Ponceau S staining (bottom panels).

### Analysis of TPST inhibition by biochemical and the protein kinase inhibitor rottlerin

The ability of fluorescent peptide substrates to report TPST1 and TPST2-directed tyrosine sulphation in a plate-based assay format allowed us to develop an enzyme screen for the analysis and discovery of small molecule TPST inhibitors. Based upon the relative ease of purification and highly stable activity towards multiple substrates, we focused our biochemical screening studies on TPST1. As detailed in Figure 5A, the TPST ligands PAP, CoA, dephospho-CoA and ATP were all able to inhibit PAPS-dependent sulphation of fluorescent CC4-tide by TPST1 *in vitro*. The IC_50_ values for inhibition ranged from low to high µM. This finding is consistent with the ability of PAP (ΔT_m_ = ~10 °C) and CoA (ΔT_m_ = ~11 °C), the two most potent inhibitors in the enzyme assay, to interact and stabilise TPST1 (and TPST2) in thermal shift assays. Interestingly, the ability of PAP (IC_50_ = 1.5 µM) and CoA (IC_50_ = 87 µM) to inhibit TPST1 sulphation activity was highly sensitive to the concentration of PAPS in the assay, with peptide sulphation in the assay increasing (less inhibition) as a function of increasing PAPS, even taking into account the increases in enzyme activity induced by high levels of PAPS (Figure 5B, C). In contrast, weak TPST1 inhibition by ATP and dephospho-CoA was largely insensitive to increases in PAPS levels in the assay, suggesting that is was likely to represent weak or non-competitive enzyme binding (Figure 5D, E).

**Figure 5.**
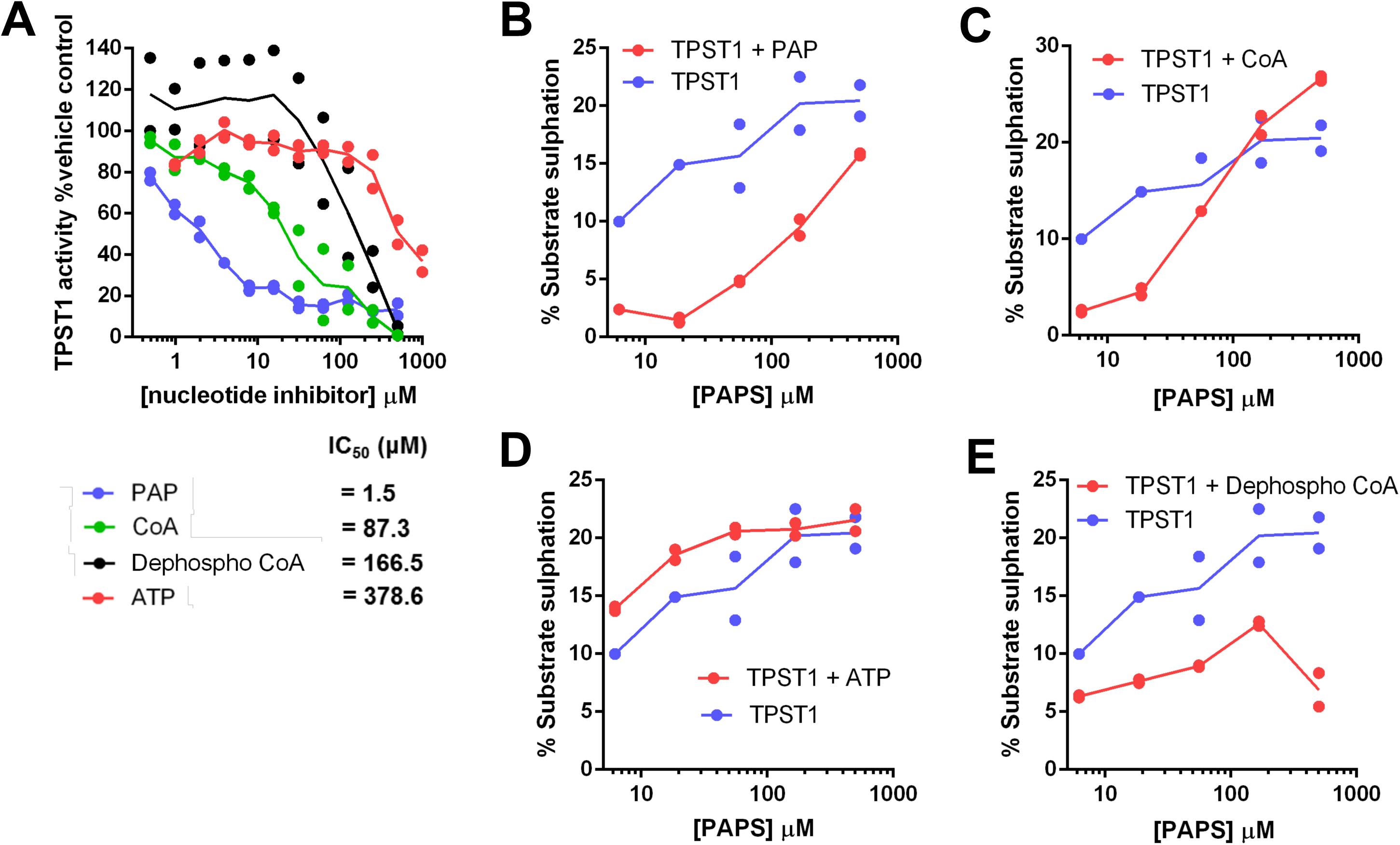
Nucleotide-dependent inhibition of TPST1 sulphotransferase activity varies with PAPS. **(A)** Dose response curves and IC_50_ values for a panel of nucleotides incubated with TPST1 in the presence of 10 μM PAPS co-factor. TPST1 activity was measured using CC4-tide and normalized to controls containing buffer alone. **(B-E)** TPST1-dependent CC4-tide sulphation was measured in the presence of increasing PAPS concentration and a fixed concentration of **(B)** 20 μM PAP, **(C)** 100 μM ATP, **(D)** 100 μM CoA or **(E)** 100 μM dephospho-CoA. All assays were performed using 0.1 μM TPST1 in the absence of MgCl_2_.

Several literature reports suggest that TPST1/2 are inhibited by nucleotide and non-nucleotide compounds *in vitro* [40, 58, 59]. Using quantitative TPST1 and 2 enzyme assays, we identified that the broad-spectrum kinase inhibitor rottlerin, which was originally described as a ‘specific’ cellular PKC inhibitor [60], but later revealed to be a non-specific protein kinase inhibitor [61], inhibited PAPS-dependent TPST1 and TPST2 with single-digit micromolar IC_50_ values (Figure 6A). We also evaluated the clinical orphan compound suramin [62] and the DNA polymerase inhibitor aurintricarboxylic acid [63] as TPST inhibitors (Figure 6C), demonstrating inhibition with low micromolar IC_50_ values, validating recent independent findings [40]. Consistently, we confirmed that rottlerin binding also induced a positive TPST1 and TPST2 thermal shift (Figure 6B), although the degree of stabilisation relative to ATP was lower than predicted, given the potent inhibitory effect of rottlerin on TPST1 and TPST2 enzyme activity *in vitro*.

**Figure 6.**
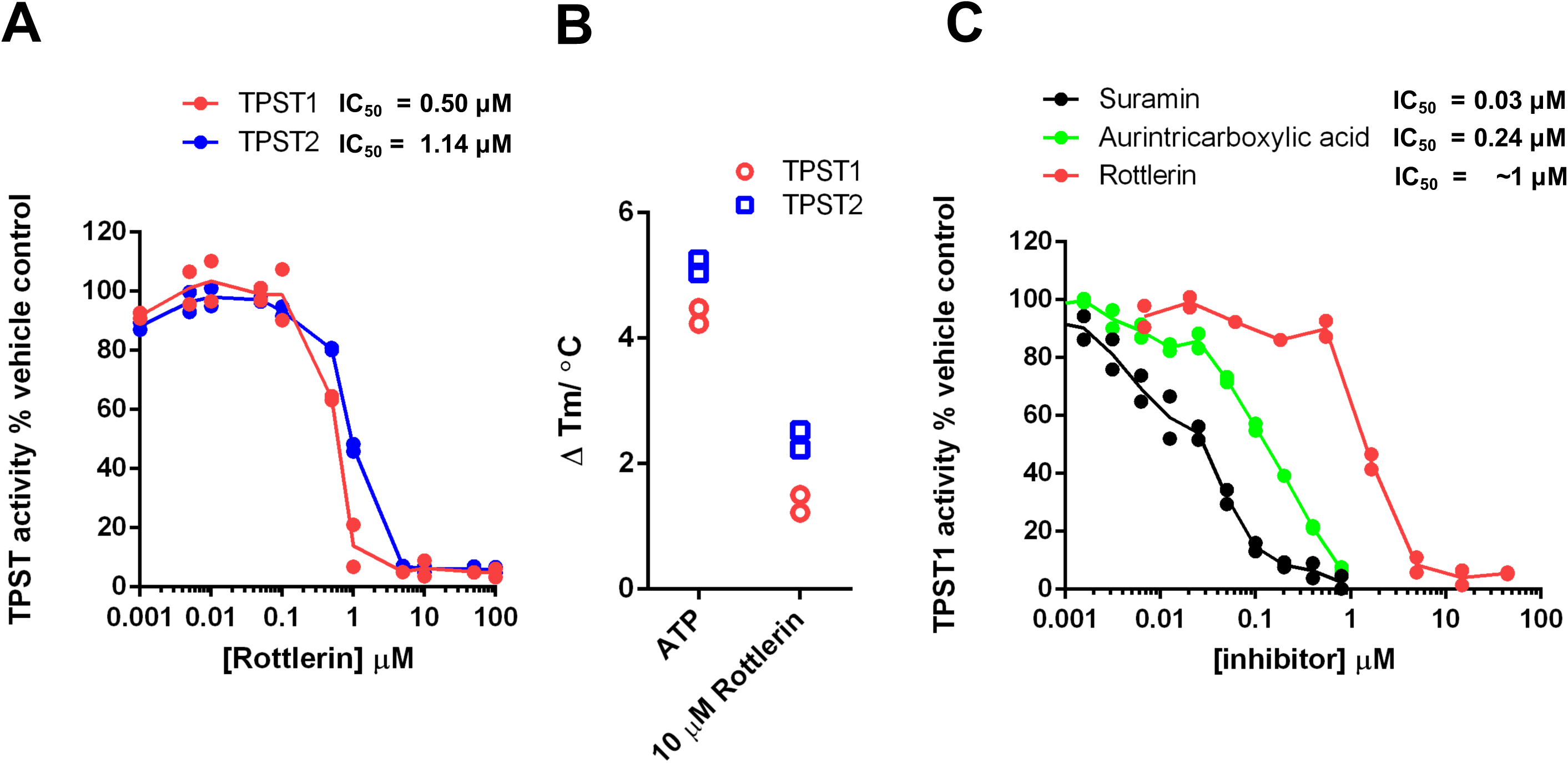
Targeting TPST sulphotransferase activity with small molecule inhibitors. **(A)** Dose response curves and calculated TPST IC_50_ values for rottlerin. TPST1 and 2 were incubated with the indicated concentration of rottlerin in the presence of 10 μM PAPS. TPST sulphotransferase activity towards CC4-tide was normalized to control reactions containing 1% (v/v) DMSO. **(B)** Thermal stability of purified TPST1 or TPST2 (5 μM) was measured in the presence of 10 μM rottlerin or 500 μM ATP as control. ∆T_m_ values were calculated by DSF as previously described. **(C)** TPST1 IC_50_ values for the previously described compounds suramin, aurintricarboxylic acid and rottlerin were calculated using TPST1 and CC4-tide in the presence of 10 μM PAPS, and activity was normalized to control reactions containing 1% (v/v) DMSO

### Multiple RAF kinase inhibitors target TPST catalytic activity *in vitro*

The provocative chemical and structural similarities between CoA, PAP, ATP and PAPS (Figure 1A), combined with the inhibitory effects of rottlerin on TPST1 and 2, raised questions about the general sensitivity of TPST enzymes to ATP-competitive kinase inhibitors. These findings prompted us to screen the open access Published Kinase Inhibitor Set (PKIS), a collection of high-quality class-annotated kinase inhibitors assembled as a starting point to discover new chemical probes for enzyme targets. The commonality of the nucleotide-binding site in huge numbers of human proteins, and shared PAPS co-factor specificity in sulphotransferases made PKIS an attractive, unbiased, resource for identifying potential new inhibitors for this family of enzymes. We took a dual-pronged approach for ligand screening, employing firstly a rapid TPST1 DSF assay and secondly a TPST1 enzyme assay. For DSF, 20 µM compound was employed for screening with 1 mM ATP as positive control, and we used a reproducible cut off value of ~ΔT_m_ ± 0.5°C in order to define a ‘hit’ (Figure 7A). The top compound found through this approach was GW406108X, and we noted strong thermal shifts in TPST1 by several compounds belonging to this indole-based kinase inhibitor class (red, Figure 7A and Supplementary Figure 4). Each ‘hit’ compound was next re-screened in a TPST1 enzyme sulphation assay at 40 µM, and the activity remaining compared to DMSO control (Figure 7B). Consistent with our DSF assay, five of the top seven TPST1 inhibitors were previously known RAF inhibitors with IC_50_ values for TPST1 in the low µM range, approximately an order of magnitude less potent than that of rottlerin (compare Figures 6C and 7C). These compounds were from two distinct types of previously described RAF inhibitors: derivatives of indole [64] or aza-stilbene [65] chemical classes. We investigated the rank order of potency for various indole compounds using GST-CC4-tide tyrosine sulphation, which was compared with rottlerin and PAP using the sulphotryosine-specific antibody (Figure 7D). Some limited Structure Activity Relationships (SARs) emerged from these initial screens, prompting us to evaluate whether our data permitted us to predict generalised TPST inhibition by other RAF inhibitors, including clinically-approved [66] and probe [67] compounds (Supplementary Figure 4). As shown in Figure 8A, duplicate assays revealed potent inhibition at a high (400 µM) concentration by many, but not all, RAF inhibitors tested, with dabrafenib, RAF-65, ZM336372, sorafenib and vemurafenib showing essentially complete TPST1 inhibition at this concentration. Titration of each compound confirmed a complex profile of inhibition, with some RAF inhibitors (e.g. vemurafenib) potentially inducing partial TPST-1 activation at lower concentrations, and then inhibiting activity at higher concentrations (Figure 8B), perhaps consistent with their complex mode of interaction with RAF, which includes promotion of dimerization and activation [68, 69]. The most compelling inhibitory data was obtained with RAF265, a phase I imidazo-benzimidazole RAF inhibitor [70, 71], for which an IC_50_ value of 6.5 µM towards TPST1 was measured, some 10-fold higher than that of rottlerin (Figure 8B). As shown in Figure 8C, both compounds exhibited dose-dependent inhibition when assayed in the presence of TPST1 and PAPS using GST-CC4-tide, and inhibition by RAF265 could be competitively decreased by increasing the concentration of PAPS in the assay, suggesting a partially competitive mode of inhibition with PAPS (Figure 8D, E).

**Figure 7.**
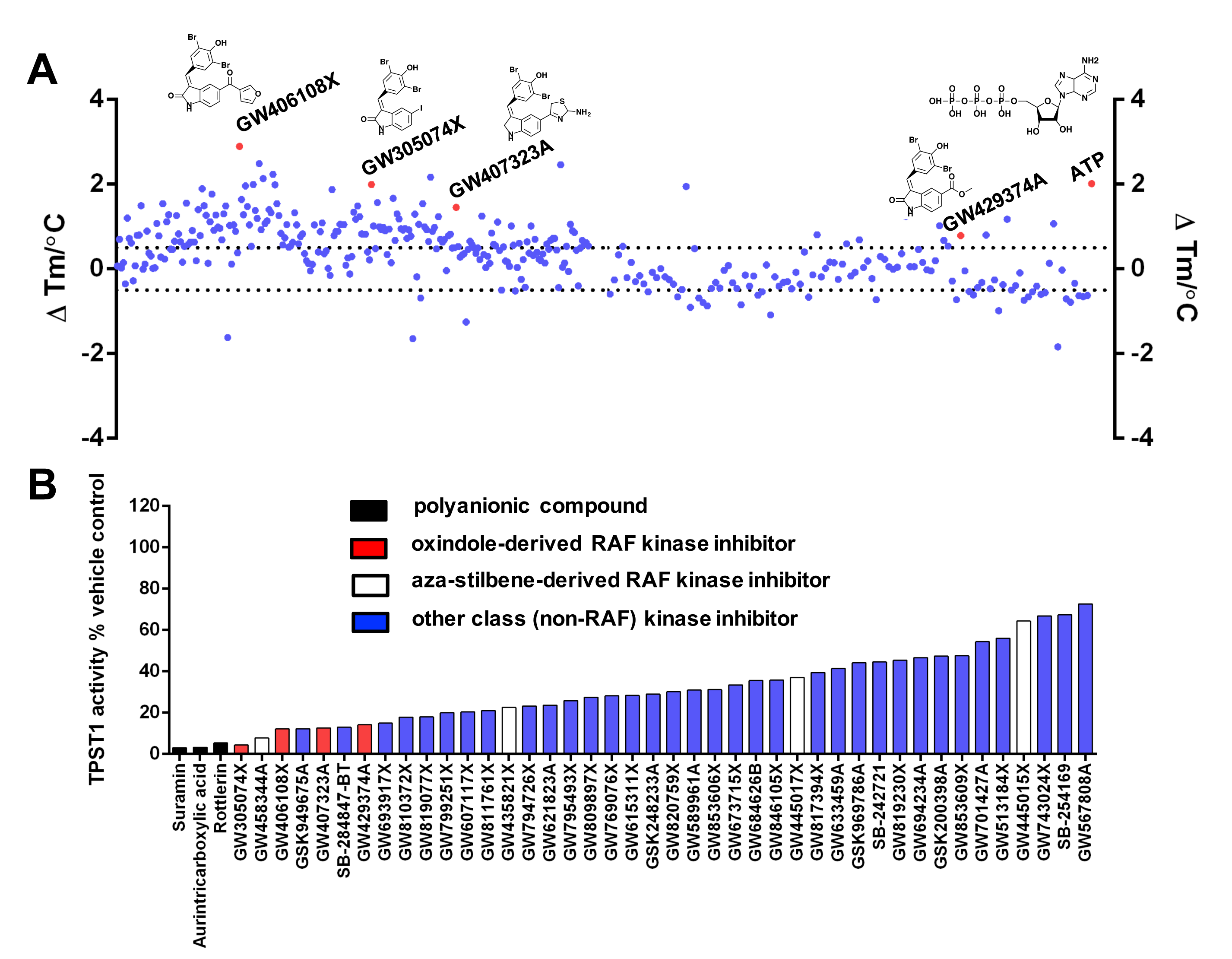
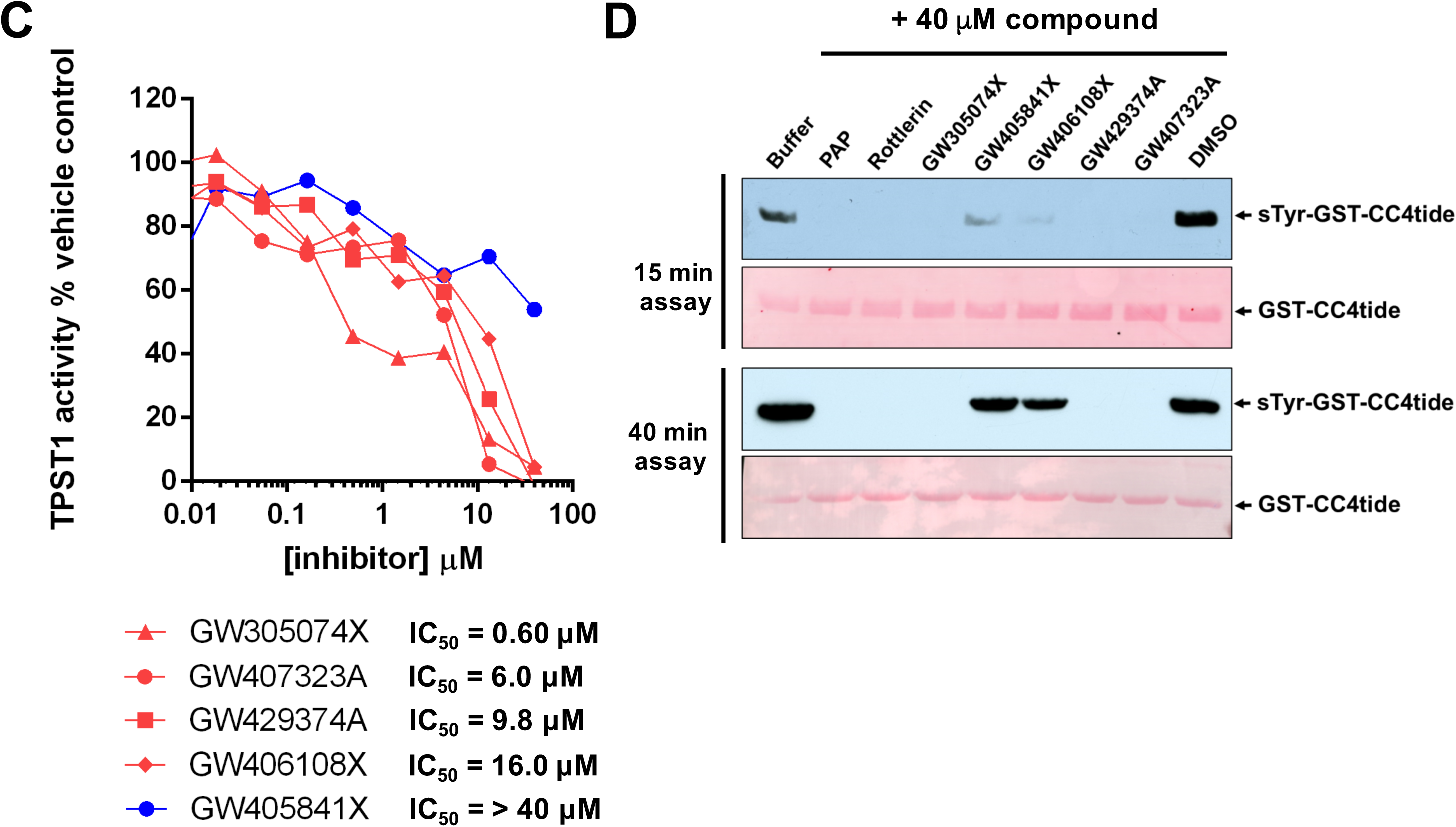
Mining the PKIS inhibitor library for TPST1 inhibitors. **(A)** Identification of PKIS small molecule ligands that alter TPST1 thermal stability. TPST1 (5 μM) was screened using PKIS compounds at final concentration of 20 μM compound and 4 % (v/v) DMSO. ∆T_m_ values were calculated by subtracting the control T_m_ value (DMSO alone, no inhibitor) from T_m_ values. Data shown is a scatter plot of the mean ∆T_m_ values from two independent DSF-based assays. **(B)** Enzymatic inhibition of TPST1 sulphotransferase activity by selected PKIS compounds. TPST1 (0.1 μM) was incubated with the appropriate PKIS compound (40 μM) in the presence of 10 μM PAPS for 30 mins at 37°C. TPST1 activity was measured using CC4-tide and normalised to 4% (v/v) DMSO control. The chemical class of inhibitor identified is colour coded. **(C)** Compound dose-response and estimation of IC_50_ values for selected chemical classes of PKIS inhibitors. TPST1 (0.1 μM) was incubated with increasing concentrations of the indicated inhibitor in the presence of 10 μM PAPS for 30 mins at 37 °C. TPST1 activity was measured using CC4-tide and normalised to DMSO controls. IC_50_ values were calculated from a single experiment, although similar results were seen in an independent experiment. **(D)** Immunoblots evaluating time-dependence of TPST1 sulphotransferase activity in the presence of a panel of PKIS or control inhibitors. 1 μg GST-CC4-tide was incubated for the appropriate time in the presence of 0.2 μg TPST1, 10 μM PAPS and a fixed concentration (40 μM) of the indicated inhibitor. After reaction termination, tyrosine sulphation was subsequently visualized using monoclonal sulphotyrosine antibody (top panel), with equal GST-CC4-tide loading confirmed by Ponceau S staining (bottom panel). GST-CC4 sulphation was performed for either 15 minutes (top panels) or 40 minutes (bottom panels).

**Figure 8.**
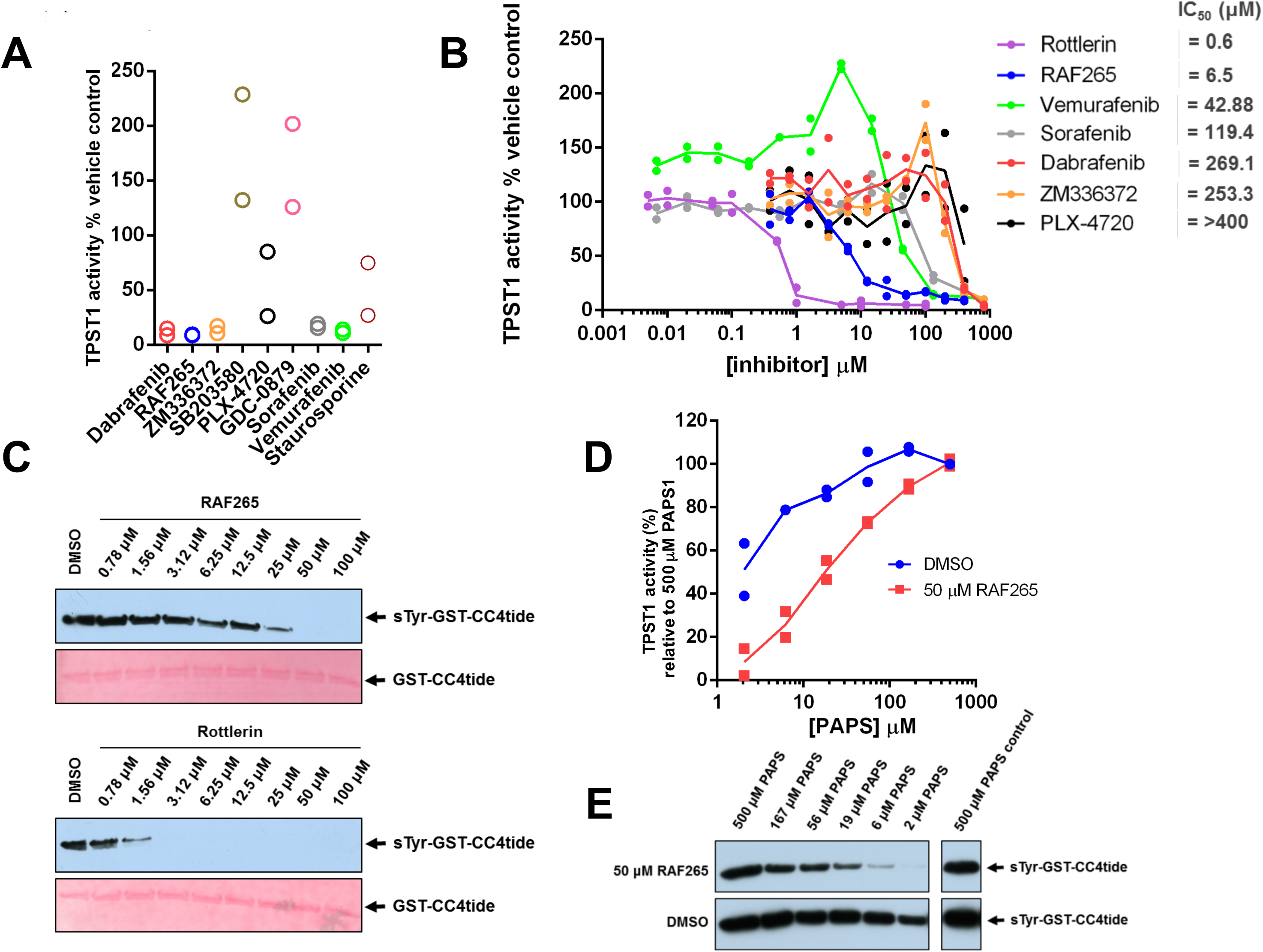
Evaluation of TPST1 inhibition by a panel of RAF kinase inhibitors. **(A)** Inhibition of TPST1 by RAF265 and other classes of RAF inhibitor. TPST1 (0.1 μM) was pre-incubated with the appropriate inhibitor (400 μM) and the assay was initiated with PAPS (10 μM). **(B)** Dose response and estimated IC_50_ values for TPST1 inhibition by RAF kinase inhibitors. TPST1 (0.1 μM) was pre-incubated with the indicated concentration of compound, and the assay initiated with PAPS (10 μM). **(C)** Immublotting of GST-CC4-tide (1 μg) sulphation by TPST1 (1 μg) in the presence of increasing concentrations of RAF265 or Rottlerin. TPST1 was pre-incubated with the indicated concentration of inhibitor, and assays performed in the presence or 10 μM PAPS for 15 mins at 20°C. **(D)** Antibody-based quantification of GST-CC4-tide sulphation by TPST1 in the presence of RAF-265 as a function of PAPS concentration. The tyrosine sulphation of GST-CC4-tide (a measure of TPST1 activity) was quantified by densitometry with IMAGE J software. Data were normalized to sulphation in the presence of 500 μM PAPS and 4% (v/v) DMSO, which represents 100% activity in the absence of the inhibitor. **(E)** A representative immunoblot corresponding to the data quantified in **(D)** is presented.

### Docking analysis of TPST ligands

In order to model the interaction of hit and control TPST1/2 ligands, including rottlerin, suramin, the sorafenib-derivative RAF265, and GW305074X with TPST1, we employed molecular docking to evaluate potential binding modes of compounds using the crystal structure of TPST1 (PDB ID: 5WRI). As shown in Figure 9A, like TPST2 [29], TPST1 possesses two adjacent docking sites in the extended catalytic region that accommodate binding of substrates, placing the tyrosine-containing substrate (left site) in proximity to the sulphate group of PAPS (right site). A docking protocol for the sulphation end-product PAP (adenosine-3’-5’-diphosphate) was developed that almost perfectly matched the crystallographic binding pose of this ligand for TPST1 (RMSD 0.30 Å, Figure 9B). By comparing experimentally-favoured configurations with those of the crystallised ligands (PAP and a CC4 peptide poised for sulphation), we were able to confidently dock rottlerin, suramin, RAF265 and GW305074X into the TPST1 active site. These compounds are predicted to make a number of stabilising interactions that help explain their ability to act as inhibitors of TPSTs *in vitro* (Figure 9C F). For example, rottlerin (C) and GW305074X (D) are predicted to occupy the peptide-binding site of TPST1, whilst suramin (E) and RAF265 (F) span both the peptide and PAPS-binding sites, consistent with the competitive loss of TPST1 inhibition by RAF265 observed as the concentration of PAPS increases in enzyme assays (Figure 8D). In contrast, suramin is predicted to form a hydrogen bond with Ser286, whilst RAF-265 forms hydrogen bonds with both Ser286 and Leu84.

**Figure 9.**
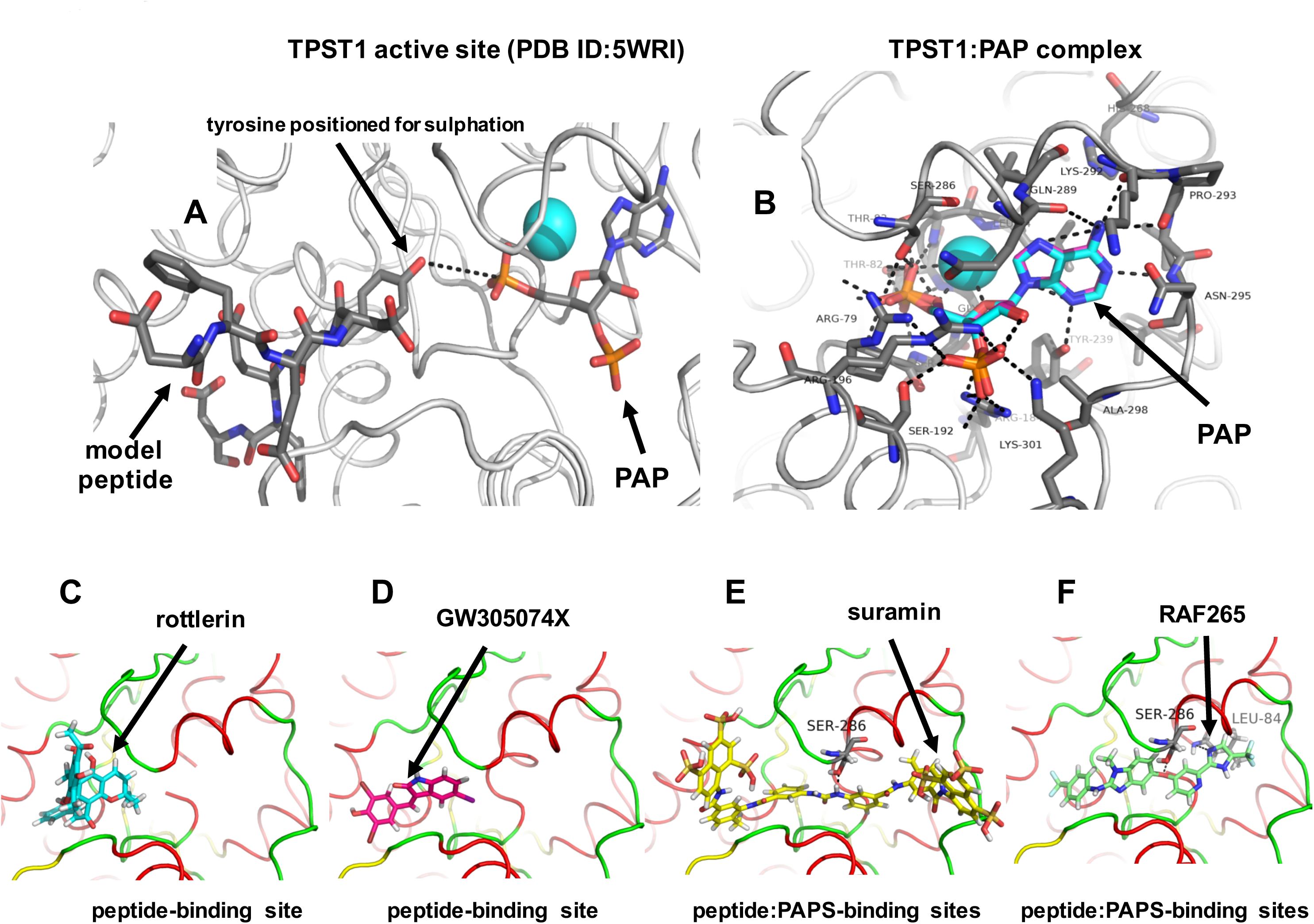
Molecular docking analysis of TPST1 with small molecule inhibitor compounds. (A) Structure of human TPST1 complexed with adenosine-3’-5’-diphosphate (PAP) and the human CC4-derived substrate EDFEDYEFD PDB ID: 5WRI (Protein rendered in grey cartoon). The inhibitory co-factor PAP (which replaces the physiological co-factor PAPS) and the co-crystallised CC4 substrate peptide are rendered as coloured sticks. Atoms are coloured grey (carbon), red (oxygen) blue (nitrogen)or cyan (oxygen of crystallographic water). Black dotted line indicates the close proximity of the tyrosyl hydroxyl group and PAP. (B) TPST1 docking poses compared. Experimentally-derived (PDB ID: 5WRI) crystallographic carbons (cyan) or our modelled docking carbons (purple) are overlayed for the inhibitory co-factor mimic PAP. TPST1 was rendered as a cartoon. PAP shown in coloured sticks. Black dotted lines indicate hydrogen bonds. Rottlerin (C), GW305074X, (D) suramin (E) or RAF265 (E) were all docked into human TPST1 (PDB ID: 5WRI), although docking solutions for each inhibitor could also be made with the very similar by employing the TPST2 catalytic domain (PDB ID: 3AP1). Proteins are depicted as cartoons with the following features: red – α helix, yellow – β sheet, green – loop. Docked molecules are coloured as sticks. Black dotted lines indicate potential hydrogen bonds.

## DISCUSSION

TPSTs catalyze protein sulphation using PAPS as the sulphate group donor, and are thought to possess structural [29] and biochemical similarities with protein tyrosine kinases relevant to both binding of synthetic substrates and an ability to modify them enzymatically *in vitro* [72]. Overlapping sulphate or phosphate modifications can potentially occur on the same tyrosine residue when appropriate acidic residues dock the substrate into the active site for covalent modification [26, 73]. Although the physiological relevance of combinatorial and competitive tyrosine modification on phosphate or sulphate (or nitrate) remains essentially unknown, bioinformatics analysis incorporating secondary structural analysis predicts that >20,000 context-dependent protein tyrosine residues are sulphated in the human proteome [30, 31, 74]. However, due to a lack of chemical tool compounds, sulphation is understudied in living organisms, often relying on ‘sledgehammer’ approaches employing non-specific reagents such as chorate or total genetic ablation [33]. The analysis of tyrosine sulphation remains ripe for both technological innovation and the discovery of new classes of sulphotransferase inhibitor [75] to promote chemical biology approaches in the field.

### DSF and sulphotransfer analysis of TPST1 and TPST2

In this paper, we report a simple and rapid method for the detection of TPST-catalysed peptide sulphation using model substrates fused to an N-terminal fluorophore. The chemical similarity between the phosphate donor ATP and PAPS, the universal sulphate donor, led us to investigate whether peptide tyrosyl sulphation could be detected using a high-throughput enzymatic procedure previously validated for phosphorylation catalysed by ATP-dependent kinases. To isolate pure, enzymatically active recombinant TPST1 and TPST2, both were expressed at high levels in bacteria, and refolded after purification from inclusion bodies using published ‘slow’ procedures suitable for structural and enzymatic analysis of TPST1 [24] and TPST2 [29]. The affinity of our TPST1 and TPST2 preparations for the PAPS co-factor was found to be almost identical to that previously reported, and we confirmed that TPST1 and TPST2 were folded and could bind to a variety of physiological and non-physiological ligands. These included sulphated PAPS and PAP, the end product of the sulphotransferase reaction (Figure 1 and Supplementary Figure 1). Protein kinases are also known bind to the reaction end product of the phosphotransferase reaction (ADP), which can act as a weak ATP-competitive inhibitor [53]. Our study also revealed that TPST1 and 2 interact with the 3’-phospho-adenosine moiety of the ligand Coenzyme A (CoA), confirming the availability of the 3’-phospho-adenosine docking region in the active site of TPSTs for unrelated ligand binding. To our knowledge, our studies are the first to employ DSF-based thermal shift assays to analyse TPST ligand binding, although these approaches are also widely used for semi-quantitative ligand binding analysis of growth factors [44, 76], protein kinase [53-56] and pseudokinase [77] domains, BH3 domains [78] and bromodomains [79].

We confirmed by DSF that TPST ligands act as competitive active-site inhibitors of peptide sulphation, creating a new impetus to develop novel screening approaches to discover TPST inhibitors. Standard biochemical assays often involve the detection of ^35^S-based substrate sulphation derived from ^35^S-labelled PAPS, and require enzymatic co-factor synthesis and time-consuming radioactive solid-phase chromatography (typically HPLC) procedures [80, 81]. In contrast, our peptide sulphation assay detects modification in real-time using a simple mobility shift assay, which is quantified by comparing the ratio of the sulphated and non-sulphated fluorescent substrates. This assay employs the EZ-Reader II platform originally developed for the rapid analysis of peptide phosphorylation, acetylation or proteolysis [48] and permits the inclusion of high concentrations of non-radioactive co-factors, substrates and ligands. The coupling of a fluorophore at the peptide N terminus, distinct from the site of tyrosine sulphation (Figure 1B) overcomes current limitations with fluorescent TPST substrates, in which the fluorophore lies adjacent to the site of sulphation. In the course of our studies, we established a high reproducibility for this assay, and exploited it to probe substrate specificity and discover new enzyme inhibitors. We also generated a substrate lacking a key Tyr residue, a dual protein kinase/TPST substrate and model TPST substrates containing acidic residues at the -1 and +1 position relative to the sulphated Tyr. These allowed us to generate CCR8 and FGF7 substrates for TPST1, which were not substrates for TPST2, and dual substrates with differential (CC4-tide) or very similar (PSGL1) sulphation kinetics for TPST1 and TPST2. Based on our initial analyses, we found that TPST2 only sulphated tyrosine-containing peptide substrates with an acidic residue in both the +1 and -1 position, whereas TPST1 dependent tyrosine sulphation only required a negative charge to be present in the -1 site (Figure 2). Future work will employ a much larger selection of peptide substrates to evaluate this preference further, with a goal of defining TPST1 and TPST2 substrate specificity *in vitro* that can be exploited to help examine the sulphoproteomic datasets emerging from cell-based studies.

### New small molecule TPST1 inhibitors

Our finding that TPST1 was inhibited at sub-micromolar concentrations by the anti-angiogenic compound suramin, which has been used clinically to treat River Blindness and African trypanasomiasis [62], and the cellular DNA polymerase inhibitor aurintricarboxylic acid [63] were intriguing, and consistent with a very recent report demonstrating inhibitory activity towards TPSTs [40]. By screening a panel of kinase inhibitors, we found that rottlerin (also known as mallotoxin) is also a low micromolar inhibitor of TPST1 and TPST2 *in vitro*. Rottlerin was originally identified as an inhibitor of PKC isozymes [60], but can also act as a sub-micromolar inhibitor of other protein kinases *in vitro* [61]. Interestingly, we discovered that all three of these compounds also have inhibitory activity towards the related PAPS-dependent oligosaccharide sulphotransferase HS2ST *in vitro* [Byrne et al., submitted alongside this manuscript], allowing us to infer that structural similarities in the PAPS or substrate-binding regions of HS2ST and TPST1/2 present a binding surface that accommodates small polyanionic compounds, which like TPST acidic peptide substrates, presumably bind through electrostatic interactions in the enzyme active site. These findings heightened the possibility that other kinase inhibitors might also be serendipitous TPST inhibitors.

To evaluate this hypothesis, we identified a number of TPST1 ligands in PKIS. Intriguingly, 4 of the top 7 hits in this screen belonged to the same benzylidene-1H-inol-2-one (oxindole) c-RAF kinase inhibitor sub-class [64]. Moreover, of the other top 30 TPST inhibitors identified (TPST1 enzyme inhibition >40% at 20 µM), GW445015X, GW445017X, and most notably, GW458344A, were all potent c-RAF inhibitors, belonging to the chemically distinct aza-stilbene chemical class [65]. We next confirmed that distinct RAF inhibitors also possess inhibitory properties towards TPST1. Interestingly, well-validated clinical RAF inhibitors, including RAF265 (IC_50_ 6.5 µM), vemurafenib (IC_50_ ~40 µM) and the much higher micromolar TPST inhibitor sorafenib (which contains the same 2-arylaminobenzimidazole chemical scaffold found in RAF265, Supplementary Figure 7), were also TPST1 inhibitors *in vitro*. These findings demonstrate that many compounds designed as RAF inhibitors also have the ability to inhibit TPST1, providing a new impetus to exploit the huge amount of RAF inhibitor design knowledge available in private and public databases for the design and testing of TPST inhibitors. In a related paper, we demonstrated cross-reactivity of rottlerin and oxindole-based (but not aza-stilbene or other RAF inhibitors) with the glycan sulphotransferases HS2ST [Byrne et al., submitted alongside this manuscript]. Interestingly, the potency of HS2ST inhibition by oxindole-based c-RAF inhibitors was some 10-fold higher than that for TPST1, and we confirmed that TPST RAF inhibitors such as RAF265 and aza-stilbenes did not inhibit HS2ST at any concentration tested. These subtle differences suggest that although inhibitor sensitivity to this class of RAF inhibitor can be shared between two distinct classes of sulphotransferase, opportunities exist for the development of both specific and potent ligands targeted more specifically towards either HS2ST or TPSTs.

## CONCLUSIONS

In order to stimulate progress in implementing chemical biology in the sulphotransferase field, careful structure-based comparison between HS2ST, TPST1/2 and RAF kinases inhibition, and analysis of a wide variety of compound chemotypes, will be required. Our docking studies with TPST1 suggest similar binding modes for both rottlerin and the potent oxindole TPST1 inhibitor GW305074X (Figure 5), whilst suramin and RAF265 feasibly interact with the extended peptide and PAPS cofactor binding sites. It will be intriguing to confirm these binding modes through structural analysis and guided enzyme mutagenesis, and to identify drug-binding site residues in sulphotransferases that dictate inhibition. This information can then be used for careful compound analysis and the generation of drug-resistant alleles for cellular analysis, using concepts developed for compound target validation in the kinase field [82-86]. In the first instance, it will also be important to evaluate whether any of the TPST ligands identified here, particularly cellular RAF inhibitors, interfere with protein tyrosine sulphation in cells, since it remains formally possible that some of the cellular phenotypes and/or clinical effects documented with these compound classes [66], might be explained in part by “off target” effects on sulphation-based biology.

Our work also raises the possibility that TPST inhibitors might be synthesised or repurposed based upon workflows previously developed for the iteration of the different families of (RAF) kinase inhibitors. Although only two TPSTs are present in multicellular eukaryotes, the development of specific inhibitors might be challenging, given the ~90% similarity within the active site, and the presence of multiple distinct PAPS-dependent sulphotransferases in vertebrate genomes. However, in this context it is useful to recall the rapid development of the kinase inhibitor field as an exemplar. Initial scepticism about the feasibility (or even, the need) to generate specific inhibitors of protein kinases has been largely overcome [87], partly through innovative synthetic approaches, but also by a deep understanding of mechanistic and structural kinase biology available within the >500 distinct members of the human kinome [35]. An appreciation that compound polypharmacology, perhaps across multiple enzyme classes, is important for driving, and predicting, both efficacy and compound side-effects for kinase inhibitors [88-90] might also be useful for the development of TPST inhibitors.

Finally, inhibitor-based interrogation of TPST-dependent tyrosine sulphation could be employed alongside MS-based sulphoproteomics [32, 91]. This could have significant impact in various areas of research by increasing our ability to chemically control and modulate tyrosine sulphation and even manipulate sulphation of specific proteins, including those implicated in, for example, infection and inflammation. Given the close parallels between tyrosine sulphation and tyrosine phosphorylation, whose rational targeting rapidly led to the clinical analysis of dozens of small molecule inhibitors [87], we suggest that a new opportunity might also soon exist to integrate the analysis of TPST with the tools of chemical biology.

## ACKNOWLEDGEMENTS

This work was funded by a BBSRC Tools and Resources Development Grant (BB/N021703/1) and a Royal Society Research Grant (to PAE), a European Commission FET-OPEN grant (ArrestAD no.737390) to DGF, PAE and DPG, North West Cancer Research (NWCR) grants CR1088 and CR1097 and a NWCR endowment (to DGF). The SGC is a registered charity (number 1097737) that receives funds from AbbVie, Bayer Pharma AG, Boehringer Ingelheim, Canada Foundation for Innovation, Eshelman Institute for Innovation, Genome Canada, Innovative Medicines Initiative (EU/EFPIA) [ULTRA-DD grant no. 115766], Janssen, Merck KGaA Darmstadt Germany, MSD, Novartis Pharma AG, Ontario Ministry of Economic Development and Innovation, Pfizer, São Paulo Research Foundation-FAPESP, Takeda, and The Wellcome Trust [106169/ZZ14/Z]

## AUTHOR CONTRIBUTIONS

PAE obtained BBSRC grant funding with DGF. PAE, DGF, DPB, YL, PN, KR, CEE and NGB designed and executed the experiments. CW, DHD and WJZ provided compound libraries, protocols and advice. PAE wrote the paper with contributions and final approval from all of the co-authors.

## ABBREVIATIONS

DSF: Differential Scanning Fluorimetry
PAP: 3’-phosphoadenosine 5’-phosphate
PAPS: 3’-phosphoadenosine 5’-phosphosulphate
PKIS: Published Kinase Inhibitor Set
RAF: Rapidly Accelerated Fibrosarcoma
TPST: Tyrosyl Protein Sulpho Transferase
TSA: Thermal Stability Assay

---

**Supplementary Figure 1.**
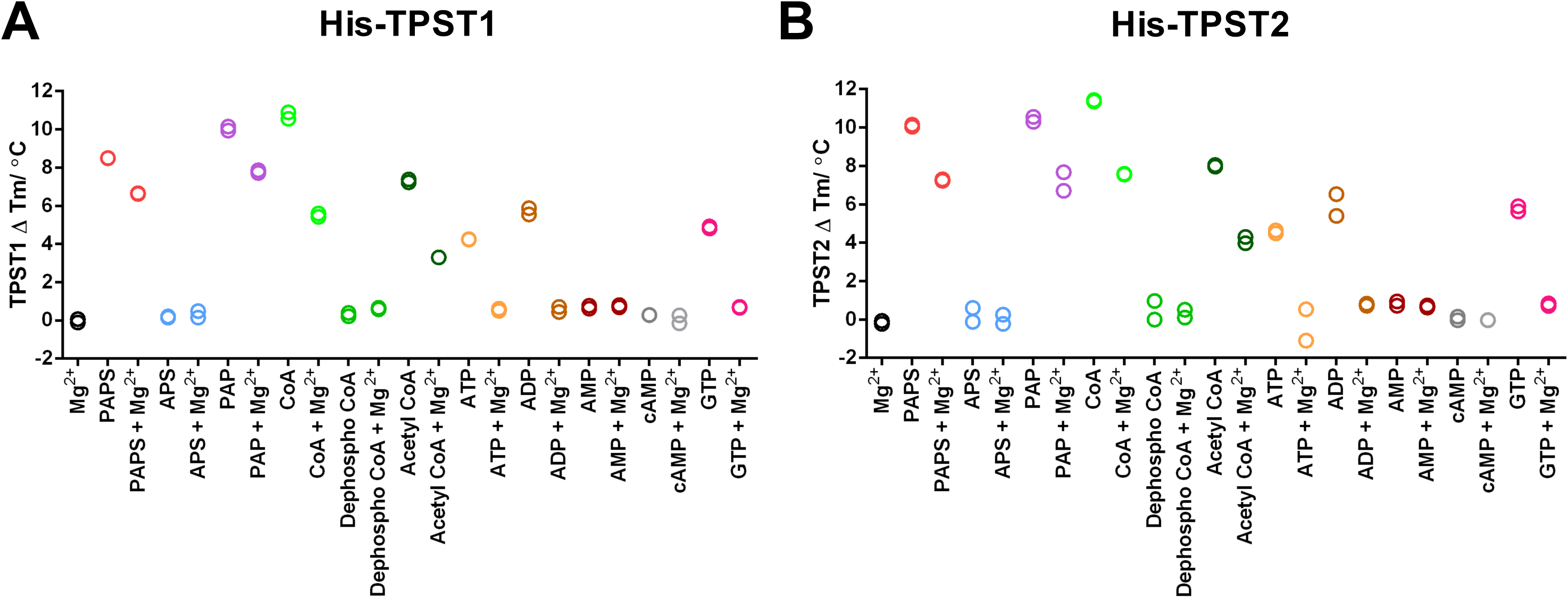
DSF-based analysis of TPST1 and TPST2 ligand interactions. Thermal stability profiles of **(A)** TPST1 or **(B)** TPST2 were measured in the presence of the indicated nucleotide. ∆T_m_ values were calculated relative to buffer controls for each TPST enzyme (5 μM) in the presence of 0.5 mM of the indicated nucleotide ± 10 mM MgCl_2_.

**Supplementary Figure 2.**
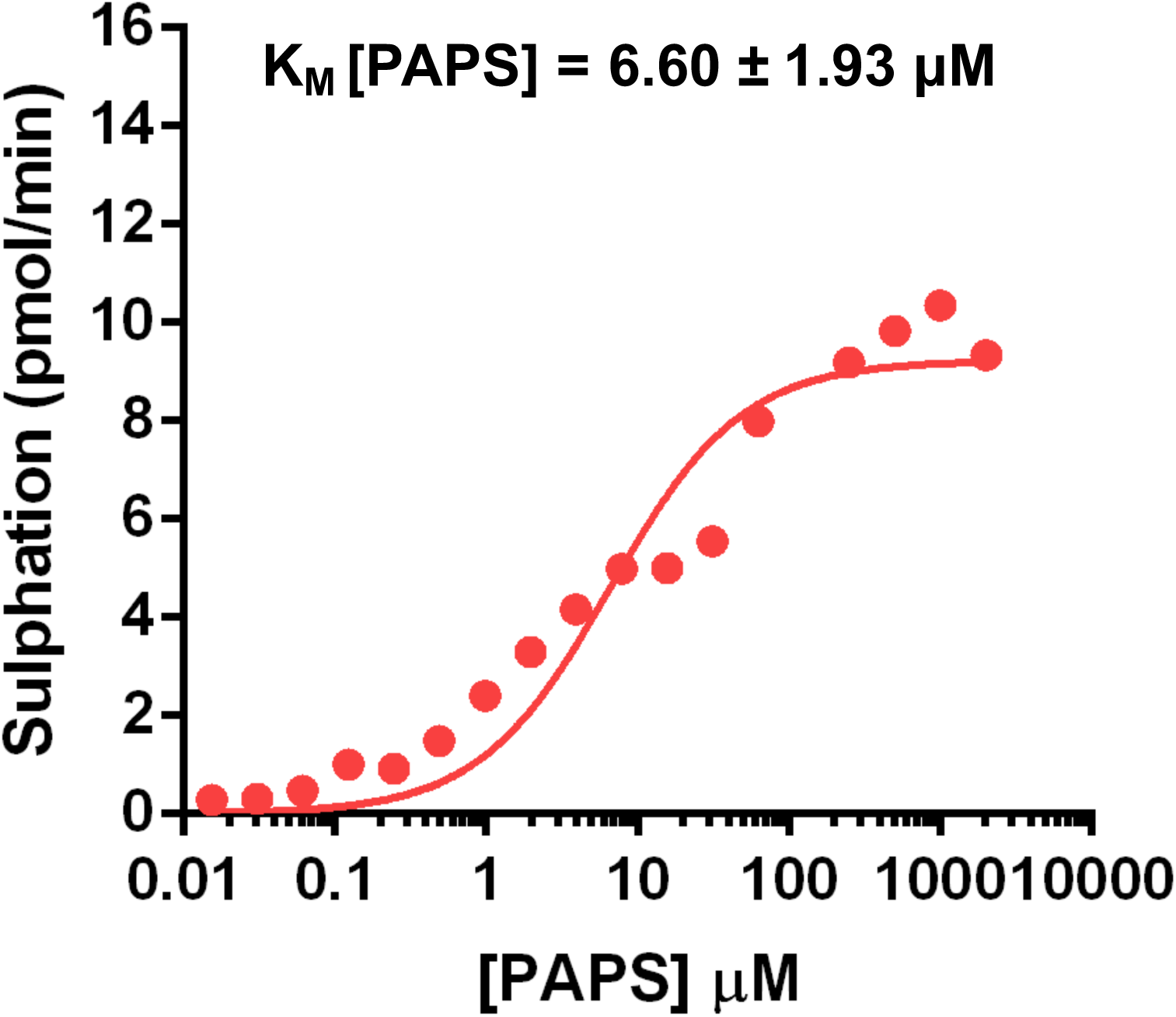
K_m_ [PAPS] determination for TPST1. Kinetic analysis of CC4-tide (2 μM) sulphation by purified TPST1 (0.1 μM) was performed in the presence of increasing concentrations of the sulphate donor PAPS. The K_m_ [PAPS] value (± standard deviation) was calculated by comparing the rate of peptide sulphation (pmoles sulphate/min) and linear regression software (GraphPad Prism) from four independent experiments.

**Supplementary Figure 3.**
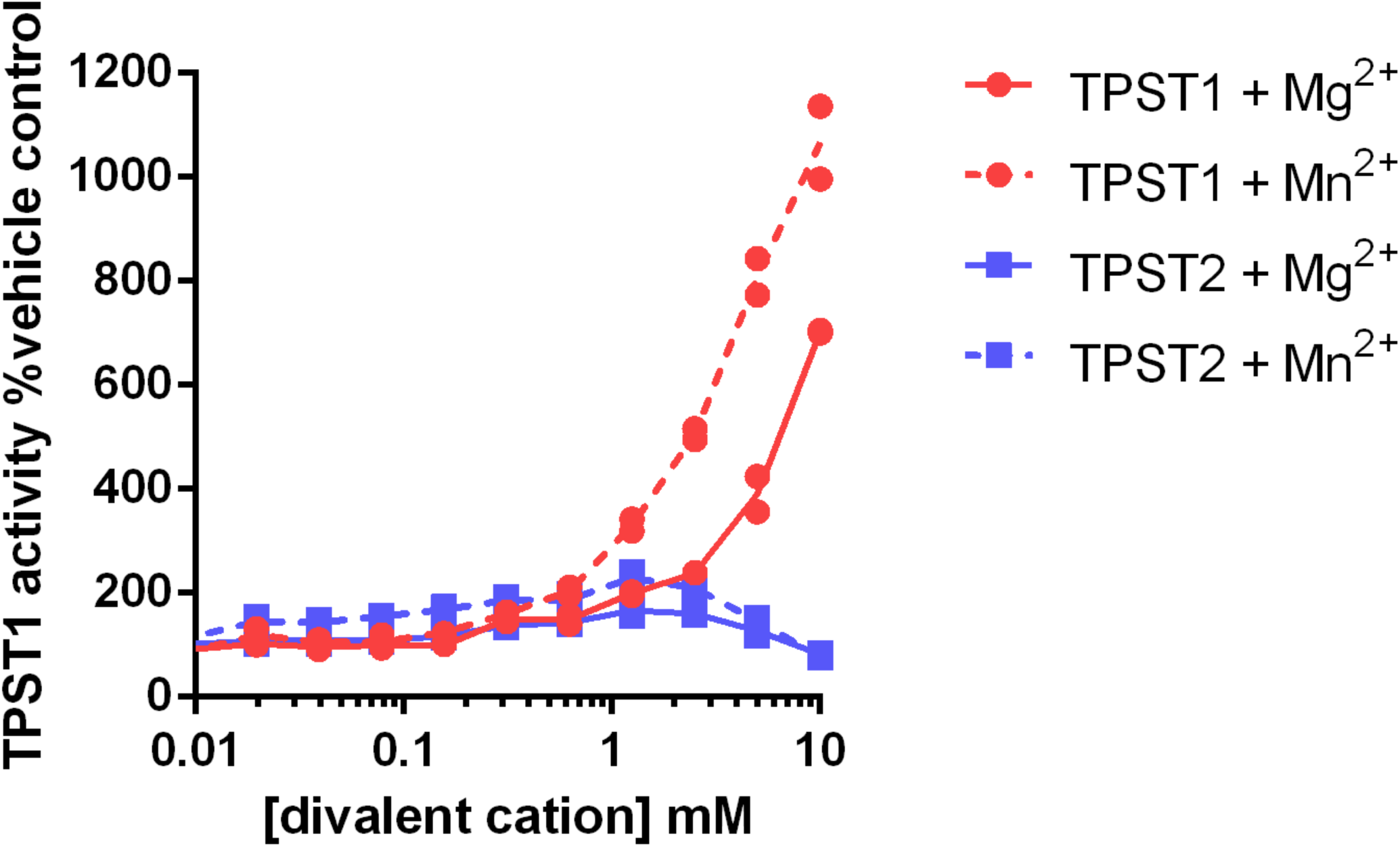
Analysis of TPST1 and TPST2 activity in the presence of selected divalent metal cations. The extent of CC4-tide sulphation was measured as a function of Mg^2+^ or Mn^2+^ ion concentration. TPST1 or TPST2 (0.1 μM) were incubated with increasing concentrations of Mg^2+^ or Mn^2+^ in the presence of 10 μM PAPS. TPST1 activity was normalised to a buffer control.

**Supplementary Figure 4.**
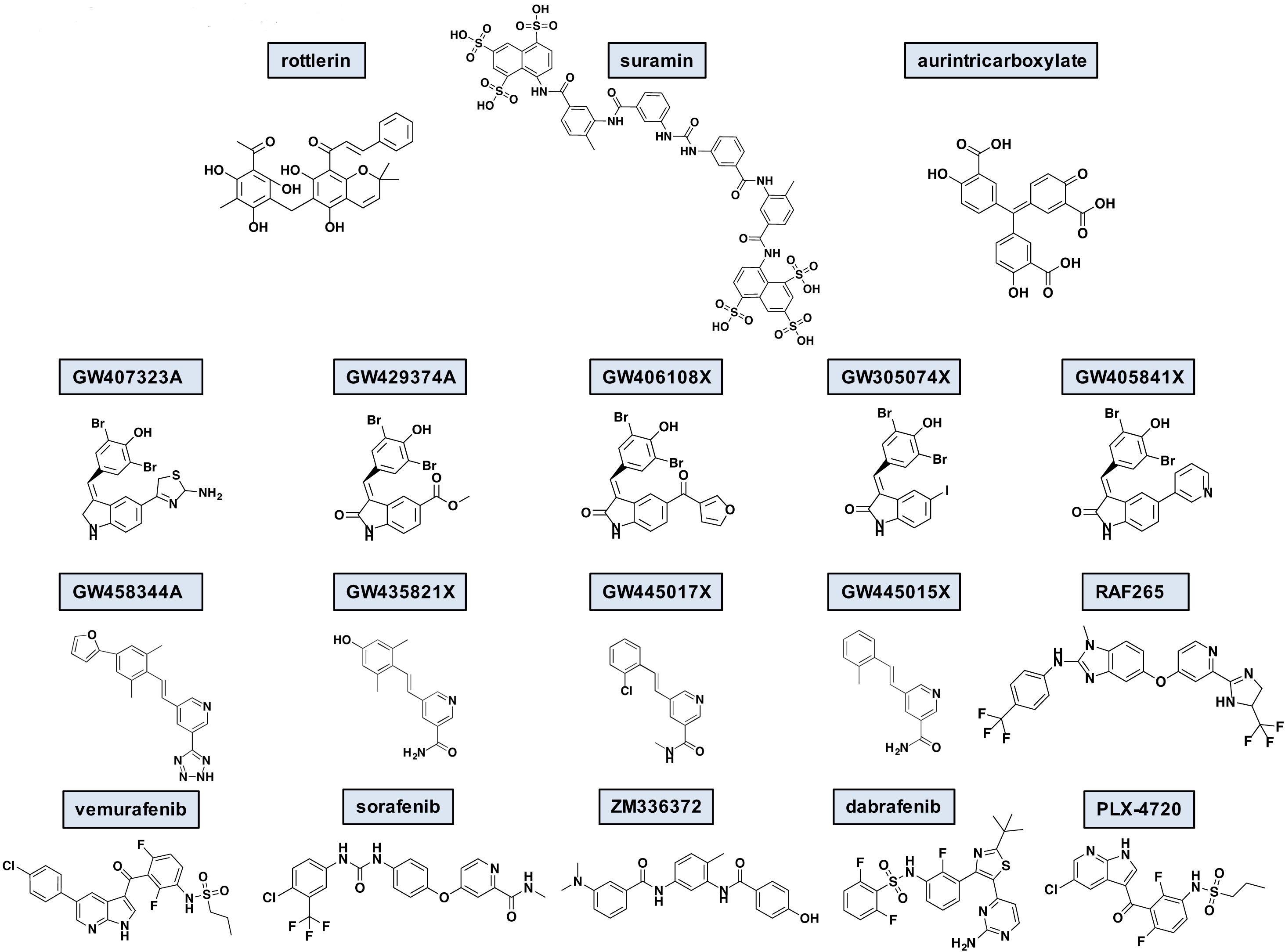
Chemical structures of TPST ligands. The chemical structures of rottlerin, suramin and aurintricarboxylic acid, a panel of TPST inhibitors discovered from PKIS and various known RAF inhibitors.

## REFERENCES

1. Hunter, T., Tyrosine phosphorylation: thirty years and counting. Current opinion in cell biology, 2009. 21(2): p. 140–6.

2. Moore, K.L., The biology and enzymology of protein tyrosine O-sulfation. The Journal of biological chemistry, 2003. 278(27): p. 24243–6.

3. Gregory, H., et al., The Antral Hormone Gastrin. Structure of Gastrin. Nature, 1964. 204: p. 931–3.

4. Ippel, J.H., et al., Structure of the tyrosine-sulfated C5a receptor N terminus in complex with chemotaxis inhibitory protein of Staphylococcus aureus. The Journal of biological chemistry, 2009. 284(18): p. 12363–72.

5. Choe, H., et al., Sulphated tyrosines mediate association of chemokines and Plasmodium vivax Duffy binding protein with the Duffy antigen/receptor for chemokines (DARC). Molecular microbiology, 2005. 55(5): p. 1413–22.

6. Veldkamp, C.T., et al., Structural basis of CXCR4 sulfotyrosine recognition by the chemokine SDF-1/CXCL12. Science signaling, 2008. 1(37): p. ra4.

7. Seibert, C., et al., Sequential tyrosine sulfation of CXCR4 by tyrosylprotein sulfotransferases. Biochemistry, 2008. 47(43): p. 11251–62.

8. Huang, C.C., et al., Structures of the CCR5 N terminus and of a tyrosine-sulfated antibody with HIV-1 gp120 and CD4. Science, 2007. 317(5846): p. 1930–4.

9. Leyte, A., et al., Sulfation of Tyr1680 of human blood coagulation factor VIII is essential for the interaction of factor VIII with von Willebrand factor. The Journal of biological chemistry, 1991. 266(2): p. 740–6.

10. Michnick, D.A., et al., Identification of individual tyrosine sulfation sites within factor VIII required for optimal activity and efficient thrombin cleavage. The Journal of biological chemistry, 1994. 269(31): p. 20095–102.

11. Cormier, E.G., et al., Specific interaction of CCR5 amino-terminal domain peptides containing sulfotyrosines with HIV-1 envelope glycoprotein gp120. Proceedings of the National Academy of Sciences of the United States of America, 2000. 97(11): p. 5762–7.

12. Farzan, M., et al., Tyrosine sulfation of the amino terminus of CCR5 facilitates HIV-1 entry. Cell, 1999. 96(5): p. 667–76.

13. Hortin, G.L., et al., Sulfation of tyrosine residues increases activity of the fourth component of complement. Proceedings of the National Academy of Sciences of the United States of America, 1989. 86(4): p. 1338–42.

14. Bundgaard, J.R., J. Vuust, and J.F. Rehfeld, Tyrosine O-sulfation promotes proteolytic processing of progastrin. The EMBO journal, 1995. 14(13): p. 3073–9.

15. Pouyani, T. and B. Seed, PSGL-1 recognition of P-selectin is controlled by a tyrosine sulfation consensus at the PSGL-1 amino terminus. Cell, 1995. 83(2): p. 333–43.

16. Bowman, K.G., et al., Identification of an N-acetylglucosamine-6-0-sulfotransferase activity specific to lymphoid tissue: an enzyme with a possible role in lymphocyte homing. Chemistry & biology, 1998. 5(8): p. 447–60.

17. Niehrs, C. and W.B. Huttner, Purification and characterization of tyrosylprotein sulfotransferase. The EMBO journal, 1990. 9(1): p. 35–42.

18. Niehrs, C., et al., Analysis of the substrate specificity of tyrosylprotein sulfotransferase using synthetic peptides. The Journal of biological chemistry, 1990. 265(15): p. 8525–32.

19. Beisswanger, R., et al., Existence of distinct tyrosylprotein sulfotransferase genes: molecular characterization of tyrosylprotein sulfotransferase-2. Proceedings of the National Academy of Sciences of the United States of America, 1998. 95(19): p. 11134–9.

20. Mishiro, E., et al., Differential enzymatic characteristics and tissue-specific expression of human TPST-1 and TPST-2. Journal of biochemistry, 2006. 140(5): p. 731–7.

21. Ouyang, Y., W.S. Lane, and K.L. Moore, Tyrosylprotein sulfotransferase: purification and molecular cloning of an enzyme that catalyzes tyrosine O-sulfation, a common posttranslational modification of eukaryotic proteins. Proceedings of the National Academy of Sciences of the United States of America, 1998. 95(6): p. 2896–901.

22. Ouyang, Y.B. and K.L. Moore, Molecular cloning and expression of human and mouse tyrosylprotein sulfotransferase-2 and a tyrosylprotein sulfotransferase homologue in Caenorhabditis elegans. The Journal of biological chemistry, 1998. 273(38): p. 24770–4.

23. Hartmann-Fatu, C., et al., Heterodimers of tyrosylprotein sulfotransferases suggest existence of a higher organization level of transferases in the membrane of the trans-Golgi apparatus. Journal of molecular biology, 2015. 427(6 Pt B): p. 1404–12.

24. Tanaka, S., et al., Structural basis for the broad substrate specificity of the human tyrosylprotein sulfotransferase-1. Scientific reports, 2017. 7(1): p. 8776.

25. Lee, R.W. and W.B. Huttner, (Glu62, Ala30, Tyr8)n serves as high-affinity substrate for tyrosylprotein sulfotransferase: a Golgi enzyme. Proceedings of the National Academy of Sciences of the United States of America, 1985. 82(18): p. 6143–7.

26. Braun, S., W.E. Raymond, and E. Racker, Synthetic tyrosine polymers as substrates and inhibitors of tyrosine-specific protein kinases. The Journal of biological chemistry, 1984. 259(4): p. 2051–4.

27. Seibert, C., et al., Tyrosine sulfation of CCR5 N-terminal peptide by tyrosylprotein sulfotransferases 1 and 2 follows a discrete pattern and temporal sequence. Proceedings of the National Academy of Sciences of the United States of America, 2002. 99(17): p. 11031–6.

28. Zhou, W., B.P. Duckworth, and R.J. Geraghty, Fluorescent peptide sensors for tyrosylprotein sulfotransferase activity. Analytical biochemistry, 2014. 461: p. 1–6.

29. Teramoto, T., et al., Crystal structure of human tyrosylprotein sulfotransferase-2 reveals the mechanism of protein tyrosine sulfation reaction. Nature communications, 2013. 4: p. 1572.

30. Monigatti, F., et al., The Sulfinator: predicting tyrosine sulfation sites in protein sequences. Bioinformatics, 2002. 18(5): p. 769–70.

31. Huang, S.Y., et al., PredSulSite: prediction of protein tyrosine sulfation sites with multiple features and analysis. Analytical biochemistry, 2012. 428(1): p. 16–23.

32. Chen, G., et al., Distinguishing Sulfotyrosine Containing Peptides from their Phosphotyrosine Counterparts Using Mass Spectrometry. Journal of the American Society for Mass Spectrometry, 2018. 29(3): p. 455–462.

33. Baeuerle, P.A. and W.B. Huttner, Chlorate--a potent inhibitor of protein sulfation in intact cells. Biochemical and biophysical research communications, 1986. 141(2): p. 870–7.

34. Hubbard, S.R., Crystal structure of the activated insulin receptor tyrosine kinase in complex with peptide substrate and ATP analog. The EMBO journal, 1997. 16(18): p. 5572–81.

35. Manning, G., et al., The protein kinase complement of the human genome. Science, 2002. 298(5600): p. 1912–34.

36. Cohen, P., Protein kinases--the major drug targets of the twenty-first century? Nature reviews. Drug discovery, 2002. 1(4): p. 309–15.

37. Kehoe, J.W., et al., Tyrosylprotein sulfotransferase inhibitors generated by combinatorial target-guided ligand assembly. Bioorganic & medicinal chemistry letters, 2002. 12(3): p. 329–32.

38. Danan, L.M., et al., Mass spectrometric kinetic analysis of human tyrosylprotein sulfotransferase-1 and -2. Journal of the American Society for Mass Spectrometry, 2008. 19(10): p. 1459–66.

39. Danan, L.M., et al., Catalytic mechanism of Golgi-resident human tyrosylprotein sulfotransferase-2: a mass spectrometry approach. Journal of the American Society for Mass Spectrometry, 2010. 21(9): p. 1633–42.

40. Zhou, W., et al., A fluorescence-based high-throughput assay to identify inhibitors of tyrosylprotein sulfotransferase activity. Biochemical and biophysical research communications, 2017. 482(4): p. 1207–1212.

41. Vargas, F., M.D. Tuong, and J.C. Schwartz, Inhibitors and dipeptide substrates for a microsomal tyrosylsulfotransferase from rat brain. Journal of enzyme inhibition, 1986. 1(2): p. 105–12.

42. Mohanty, S., et al., Hydrophobic Core Variations Provide a Structural Framework for Tyrosine Kinase Evolution and Functional Specialization. PLoS genetics, 2016. 12(2): p. e1005885.

43. Seibert, C., et al., Preparation and analysis of N-terminal chemokine receptor sulfopeptides using tyrosylprotein sulfotransferase enzymes, in Methods in enzymology 2016, Elsevier. p. 357–388.

44. Sun, C., et al., HaloTag is an effective expression and solubilisation fusion partner for a range of fibroblast growth factors. PeerJ, 2015. 3: p. e1060.

45. McSkimming, D.I., et al., KinView: a visual comparative sequence analysis tool for integrated kinome research. Molecular bioSystems, 2016. 12(12): p. 3651–3665.

46. Murphy, J.M., et al., A robust methodology to subclassify pseudokinases based on their nucleotide-binding properties. The Biochemical journal, 2014. 457(2): p. 323–34.

47. Murphy, J.M., et al., A robust methodology to subclassify pseudokinases based on their nucleotide-binding properties. Biochemical Journal, 2014. 457(2): p. 323–334.

48. Blackwell, L.J., et al., High-throughput screening of the cyclic AMP-dependent protein kinase (PKA) using the caliper microfluidic platform. Methods in molecular biology, 2009. 565: p. 225–37.

49. Elkins, J.M., et al., Comprehensive characterization of the Published Kinase Inhibitor Set. Nature biotechnology, 2016. 34(1): p. 95–103.

50. Drewry, D.H., et al., Progress towards a public chemogenomic set for protein kinases and a call for contributions. PloS one, 2017. 12(8): p. e0181585.

51. Jones, G., et al., Development and validation of a genetic algorithm for flexible docking. Journal of molecular biology, 1997. 267(3): p. 727–48.

52. Korb, O., T. Stutzle, and T.E. Exner, Empirical scoring functions for advanced protein-ligand docking with PLANTS. Journal of chemical information and modeling, 2009. 49(1): p. 84–96.

53. Byrne, D.P., et al., cAMP-dependent protein kinase (PKA) complexes probed by complementary differential scanning fluorimetry and ion mobility-mass spectrometry. The Biochemical journal, 2016. 473(19): p. 3159–75.

54. Rudolf, A.F., et al., A comparison of protein kinases inhibitor screening methods using both enzymatic activity and binding affinity determination. PloS one, 2014. 9(6): p. e98800.

55. Dodson, C.A., et al., A kinetic test characterizes kinase intramolecular and intermolecular autophosphorylation mechanisms. Science signaling, 2013. 6(282): p. ra54.

56. Caron, D., et al., Mitotic phosphotyrosine network analysis reveals that tyrosine phosphorylation regulates Polo-like kinase 1 (PLK1). Science signaling, 2016. 9(458): p. rs14.

57. Hsu, Y.R., et al., Human keratinocyte growth factor recombinantly expressed in Chinese hamster ovary cells: isolation of isoforms and characterization of post-translational modifications. Protein expression and purification, 1998. 12(2): p. 189–200.

58. Esko, J.D., C. Bertozzi, and R.L. Schnaar, Chemical Tools for Inhibiting Glycosylation, in Essentials of Glycobiology, rd, et al., Editors. 2015: Cold Spring Harbor (NY). p. 701–712.

59. Armstrong, J.I. and C.R. Bertozzi, Sulfotransferases as targets for therapeutic intervention. Current opinion in drug discovery & development, 2000. 3(5): p. 502–15.

60. Gschwendt, M., et al., Rottlerin, a novel protein kinase inhibitor. Biochemical and biophysical research communications, 1994. 199(1): p. 93–8.

61. Davies, S.P., et al., Specificity and mechanism of action of some commonly used protein kinase inhibitors. The Biochemical journal, 2000. 351(Pt 1): p. 95–105.

62. McGeary, R.P., et al., Suramin: clinical uses and structure-activity relationships. Mini reviews in medicinal chemistry, 2008. 8(13): p. 1384–94.

63. Givens, J.F. and K.F. Manly, Inhibition of RNA-directed DNA polymerase by aurintricarboxylic acid. Nucleic acids research, 1976. 3(2): p. 405–18.

64. Lackey, K., et al., The discovery of potent cRaf1 kinase inhibitors. Bioorganic & medicinal chemistry letters, 2000. 10(3): p. 223–6.

65. McDonald, O., et al., Aza-stilbenes as potent and selective c-RAF inhibitors. Bioorganic & medicinal chemistry letters, 2006. 16(20): p. 5378–83.

66. Karoulia, Z., E. Gavathiotis, and P.I. Poulikakos, New perspectives for targeting RAF kinase in human cancer. Nature reviews. Cancer, 2017. 17(11): p. 676–691.

67. Arrowsmith, C.H., et al., The promise and peril of chemical probes. Nature chemical biology, 2015. 11(8): p. 536–41.

68. Hall-Jackson, C.A., et al., Paradoxical activation of Raf by a novel Raf inhibitor. Chemistry & biology, 1999. 6(8): p. 559–68.

69. Poulikakos, P.I., et al., RAF inhibitor resistance is mediated by dimerization of aberrantly spliced BRAF(V600E). Nature, 2011. 480(7377): p. 387–90.

70. Williams, T.E., et al., Discovery of RAF265: A Potent mut-B-RAF Inhibitor for the Treatment of Metastatic Melanoma. ACS medicinal chemistry letters, 2015. 6(9): p. 961–5.

71. Izar, B., et al., A first-in-human phase I, multicenter, open-label, dose-escalation study of the oral RAF/VEGFR-2 inhibitor (RAF265) in locally advanced or metastatic melanoma independent from BRAF mutation status. Cancer medicine, 2017. 6(8): p. 1904–1914.

72. Hunter, T., Synthetic peptide substrates for a tyrosine protein kinase. The Journal of biological chemistry, 1982. 257(9): p. 4843–8.

73. Baldwin, G.S., J. Knesel, and J.M. Monckton, Phosphorylation of gastrin-17 by epidermal growth factor-stimulated tyrosine kinase. Nature, 1983. 301(5899): p. 435–7.

74. Yang, Y.S., et al., Tyrosine sulfation as a protein post-translational modification. Molecules, 2015. 20(2): p. 2138–64.

75. Rath, V.L., D. Verdugo, and S. Hemmerich, Sulfotransferase structural biology and inhibitor discovery. Drug discovery today, 2004. 9(23): p. 1003–11.

76. Li, Y., et al., Heparin binding preference and structures in the fibroblast growth factor family parallel their evolutionary diversification. Open biology, 2016. 6(3).

77. Bailey, F.P., et al., The Tribbles 2 (TRB2) pseudokinase binds to ATP and autophosphorylates in a metal-independent manner. The Biochemical journal, 2015. 467(1): p. 47–62.

78. Milani, M., et al., DRP-1 is required for BH3 mimetic-mediated mitochondrial fragmentation and apoptosis. Cell death & disease, 2017. 8(1): p. e2552.

79. Hay, D.A., et al., Discovery and optimization of small-molecule ligands for the CBP/p300 bromodomains. Journal of the American Chemical Society, 2014. 136(26): p. 9308–19.

80. Bourdineaud, J.P., et al., Enzymatic radiolabelling to a high specific activity of legume lipooligosaccharidic nodulation factors from Rhizobium meliloti. The Biochemical journal, 1995. 306 (Pt 1): p. 259–64.

81. Armstrong, J.I., et al., Discovery of Carbohydrate Sulfotransferase Inhibitors from a Kinase-Directed Library. Angewandte Chemie, 2000. 39(7): p. 1303–1306.

82. Eyers, P.A., et al., Use of a drug-resistant mutant of stress-activated protein kinase 2a/p38 to validate the in vivo specificity of SB 203580. FEBS letters, 1999. 451(2): p. 191–6.

83. Scutt, P.J., et al., Discovery and exploitation of inhibitor-resistant aurora and polo kinase mutants for the analysis of mitotic networks. The Journal of biological chemistry, 2009. 284(23): p. 15880–93.

84. Sloane, D.A., et al., Drug-resistant aurora A mutants for cellular target validation of the small molecule kinase inhibitors MLN8054 and MLN8237. ACS chemical biology, 2010. 5(6): p. 563–76.

85. Bailey, F.P., V.I. Andreev, and P.A. Eyers, The resistance tetrad: amino acid hotspots for kinome-wide exploitation of drug-resistant protein kinase alleles. Methods in enzymology, 2014. 548: p. 117–46.

86. Bury, L., et al., Plk4 and Aurora A cooperate in the initiation of acentriolar spindle assembly in mammalian oocytes. The Journal of cell biology, 2017. 216(11): p. 3571–3590.

87. Fabbro, D., 25 years of small molecular weight kinase inhibitors: potentials and limitations. Molecular pharmacology, 2015. 87(5): p. 766–75.

88. Ferguson, F.M. and N.S. Gray, Kinase inhibitors: the road ahead. Nature reviews. Drug discovery, 2018.

89. Dar, A.C., et al., Chemical genetic discovery of targets and anti-targets for cancer polypharmacology. Nature, 2012. 486(7401): p. 80–4.

90. Klaeger, S., et al., The target landscape of clinical kinase drugs. Science, 2017. 358(6367).

91. Huang, B.Y., et al., High-Throughput Screening of Sulfated Proteins by Using a Genome-Wide Proteome Microarray and Protein Tyrosine Sulfation System. Analytical chemistry, 2017. 89(6): p. 3278–3284.

